# A rooted phylogeny resolves early bacterial evolution

**DOI:** 10.1101/2020.07.15.205187

**Authors:** Gareth A. Coleman, Adrián A. Davín, Tara Mahendrarajah, Anja Spang, Philip Hugenholtz, Gergely J. Szöllősi, Tom A. Williams

**Author notes:** Contributed equally to this paper (Gareth A. Coleman, Adrián A. Davín).

## Abstract

Bacteria are the most abundant and metabolically diverse cellular lifeforms on Earth. A rooted bacterial phylogeny provides a framework to interpret this diversity and to understand the nature of early life. Inferring the position of the bacterial root is complicated by incomplete taxon sampling and the long branch to the archaeal outgroup. To circumvent these limitations, we model bacterial genome evolution at the level of gene duplication, transfer and loss events, allowing outgroup-free inference of the root^1^. We infer a rooted bacterial tree on which 68% of gene transmission events are vertical. Our analyses reveal a basal split between Terrabacteria and Gracilicutes, which together encompass almost all known bacterial diversity. However, the position of one phylum, Fusobacteriota, could not be resolved in relation to these two major clades. In contrast to recent proposals, our analyses strongly reject a root between the Candidate Phyla Radiation (CPR) and all other Bacteria. Instead, we find that the CPR is a sister lineage to the Chloroflexota within the Terrabacteria. We predict that the last bacterial common ancestor was a free-living flagellated, rod-shaped cell featuring a double membrane with a lipopolysaccharide outer layer, a Type III CRISPR-Cas system, Type IV pili, and the ability to sense and respond via chemotaxis.

---

Rooting deep radiations is among the greatest challenges in phylogenomics, and there is no consensus on the root of the bacterial tree. Based on evidence^2–5^ that the root of the entire tree of life lies between Bacteria and Archaea, early analyses using an archaeal outgroup placed the bacterial root near Aquificales/Thermotogales^6,7^ or Planctomycetes^8^. Alternative approaches, including analyses of gene flows and polarisation of changes in multimeric protein complexes and other complex characters^9^, have instead suggested roots within the monoderm (single-membrane) Bacteria^10^, or between Chloroflexi and all other cellular life^9^. The development of techniques for sequencing microbes directly from environmental samples, without the need for laboratory cultivation, has greatly expanded the genomic representation of natural prokaryotic diversity^11–14^. Recent phylogenomic analyses of that expanded diversity have placed the bacterial root between one of these new groups, the Candidate Phyla Radiation (CPR; also known as Patescibacteria^15,16^) and all other Bacteria^11,16,17^. The CPR comprises lineages that are characterised by small cells and genomes and are suggested to have predominantly symbiotic or parasitic lifestyles, but much remains to be learned about their ecology and physiology^15,17–19^. If correct, the early divergence of CPR has important implications for our understanding of the earliest period of cellular evolution. Taken together with evidence that the root of the archaeal domain lies between the reduced and predominantly host-associated DPANN superphylum and the rest of Archaea^1,20^, the CPR root would imply that streamlined, metabolically minimalist prokaryotes have co-existed with the more familiar, self-sufficient lineages throughout the history of cellular life^19^.

Improved taxon sampling can help to resolve difficult phylogenetic problems^21,22^, and the enormous quantity and diversity of genome data now available presents an unprecedented opportunity to resolve long-standing questions about the origins and diversification of Bacteria. But deep phylogenetic divergences are difficult to resolve, both because the phylogenetic signal for deep relationships is overwritten by new changes through time, and also because the process of sequence evolution is more complex than the best-fitting models currently available. In particular, variation in nucleotide or amino acid composition across the sites of the alignment and the branches of the tree can induce long branch attraction (LBA) artifacts in which deep-branching, fast-evolving, poorly-sampled or compositionally biased lineages group together irrespective of their evolutionary history^23^. These issues are widely appreciated^11^ but are challenging to adequately address, particularly when sequences from thousands of taxa^11,13,14,16,17^ are used to estimate trees of global prokaryotic diversity, which precludes the use of the best available phylogenetic methods.

## An unrooted phylogeny of Bacteria

We investigated the topology and root of the bacterial domain on representative subsamples of known bacterial diversity - including uncultured taxa - using the best-fitting evolutionary models. Our focal analysis consisted of 265 bacterial genomes sampled evenly from across the diversity present in the Genome Taxonomy Database (GTDB)^13^; we also performed an additional analysis in which representative taxa were chosen using a different, GTDB-independent approach, to evaluate the effect of taxon sampling on our inferences (see Supplementary Methods). The OMA method^24^ was combined with manual curation of initial single-gene trees to identify 63 phylogenetic markers (Extended Data Table 1) that our analyses suggest have evolved vertically within Bacteria. We inferred an unrooted species tree from a concatenation of these 63 markers using the LG+C60+R8+F model in IQ-Tree 1.6.10 (**Figure 1**), which was chosen as the best-fitting model using the Bayesian Information Criterion. We obtained highly congruent trees when removing 20-80% of the most compositionally heterogeneous sites from the alignment (Extended Data Figure 1), suggesting that the key features of the topology are not composition-driven LBA artifacts. All trees were consistent with the GTDB taxonomy, with all widely accepted phyla being resolved as monophyletic lineages, including the proposal that the Tenericutes branch within the Firmicutes^25^. Higher-level associations of phyla were also resolved, notably PVC^26^, FCB^27^, Cyanobacteria-Magulisbacteria^28^, Chloroflexota-Dormibacterota^29^ and the CPR^15^. The largest stable groups in the unrooted tree were the Gracilicutes^9^, comprising the majority of diderm lineages; and the Terrabacteria^30^, which comprise monoderm and atypical monoderm lineages, and which in our analyses include the CPR. The position of Fusobacteriota was unstable in the compositionally-stripped trees, either branching as in the focal tree (40%, 60%, 80% of most compositionally heterogeneous site removed) or with Deinococcus-Synergistetes-Thermotogae (20% of sites removed).

**Figure 1:**
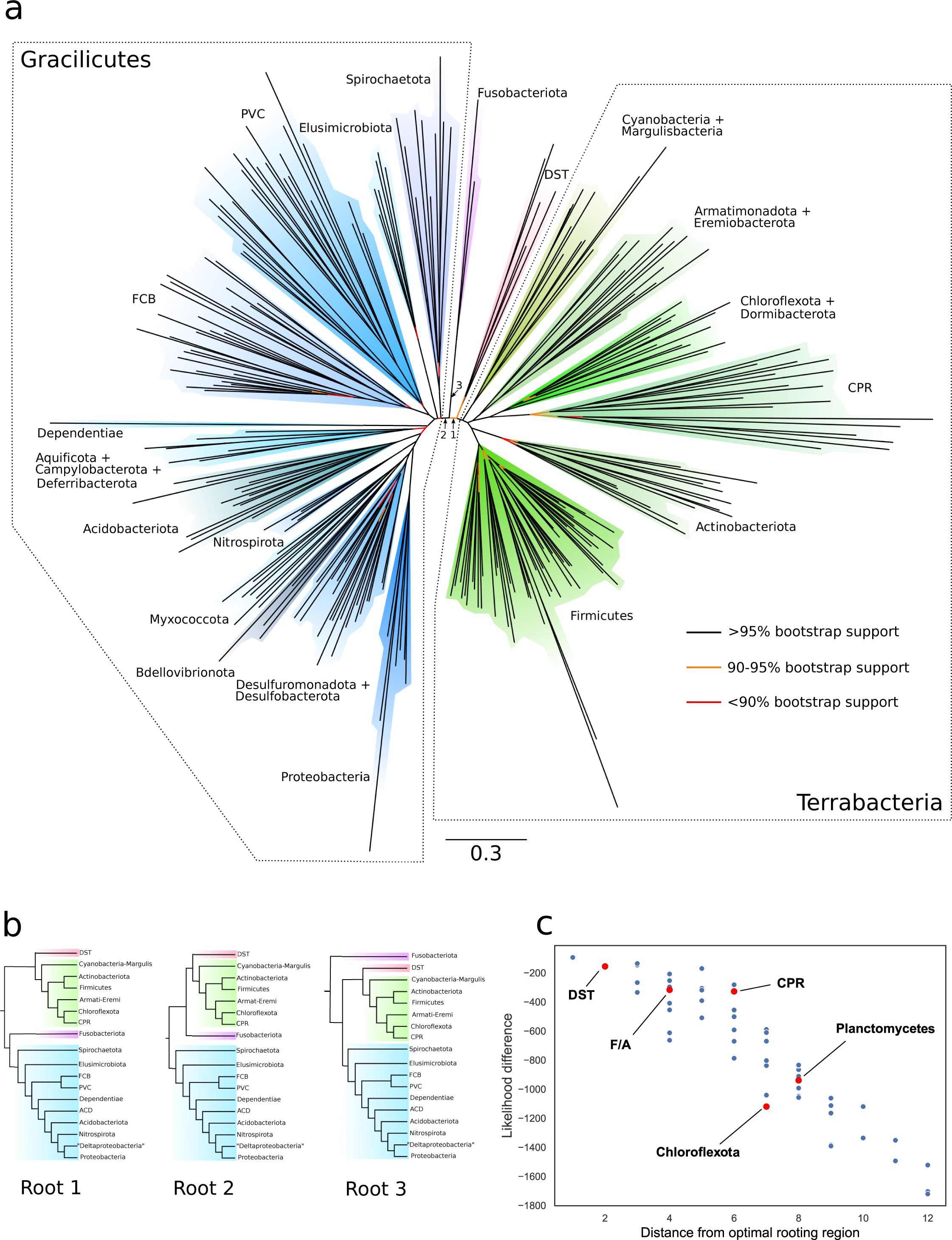
A rooted phylogeny of Bacteria. (a) We used gene tree-species tree reconciliation to infer the root of the bacterial tree. The unrooted phylogeny was inferred from a concatenation of 63 marker genes under the best-fitting LG+C60+R8+F model, which accounts for site-heterogeneity in the substitution process and uses a mixture of 8 substitution rates estimated from the data to model across-site evolutionary rate variation. Branches are coloured according to bootstrap support value. The root falls between two major clades of Bacteria, the Gracilicutes and the Terrabacteria, on one of three statistically equivalent adjacent branches indicated by arrows, shown as rooted trees in (b). All tested alternative roots were rejected (Extended Data Table 3, Supplementary 1) with likelihoods decreasing as a function of distance from the root region, as shown in (c). Previously proposed root positions, including the CPR root, are highlighted in red. FCB are the Fibrobacterota, Chlorobiota, Bacteroidota, and related lineages; PVC are the Planctomycetota, Verrucomicrobiota, Chlamydiota, and related lineages; DST are the Deinococcota, the Synergistota, and Thermotogota; ACD are Aquificota, Campylobacterota, and Deferribacterota; FA are Firmicutes and Actinobacteriota.

## Rooting the bacterial tree

The standard approach to rooting is to include an outgroup in the analysis, and all published bacterial phylogenies in which CPR form a basal lineage^11,16,17^ have made use of an archaeal outgroup. Outgroup rooting on the bacterial tree, however, has three serious limitations. First, interpretation of the results requires the assumption that the root of the tree of life lies between Bacteria and Archaea. While this is certainly the consensus view, the available evidence is limited and difficult to interpret^2–5,31^, and alternative hypotheses in which the universal root is placed within Bacteria have been proposed on the basis of indels^32,33^ or the analysis of slow-evolving characters^9^. Second, the long branch leading to the archaeal outgroup has the potential to distort within-Bacteria relationships because of LBA. Third, joint analyses of Archaea and Bacteria are based on the smaller number of genes that are widely conserved and have evolved vertically since the divergence of the two lineages, and sequence alignment is more difficult because of the great evolutionary distance between the domains.

We began by evaluating the performance of outgroup rooting on the bacterial tree using 143 Archaea and a shared subset of 30 of our phylogenetic markers (Extended Data Table 1). Using this archaeal outgroup, the ML phylogeny under the best-fitting model (LG+C60+R8+F, which accounts for site-heterogeneity in the substitution process) placed the bacterial root between a clade comprising Cyanobacteria+Margulisbacteria, CPR+Chloroflexi+Dormibacteria on one side of the root, and all other taxa on the other (Extended Data Figure 2). However, bootstrap support for this root, and indeed many other deep branches in both the bacterial and archaeal subtrees was low (50-80%). We therefore used approximately-unbiased (AU) tests^34^ to determine whether a range of published alternative rooting hypotheses (Extended Data Table 2) could be rejected, given the model and data. The AU test asks whether the optimal trees that are consistent with these other hypotheses have a significantly worse likelihood score than the maximum likelihood tree. In this case, the likelihoods of all tested trees were statistically indistinguishable (AU > 0.05, Extended Data Table 2). This indicates that outgroup rooting cannot resolve the bacterial root on this alignment of 30 conserved genes.

**Figure 2:**
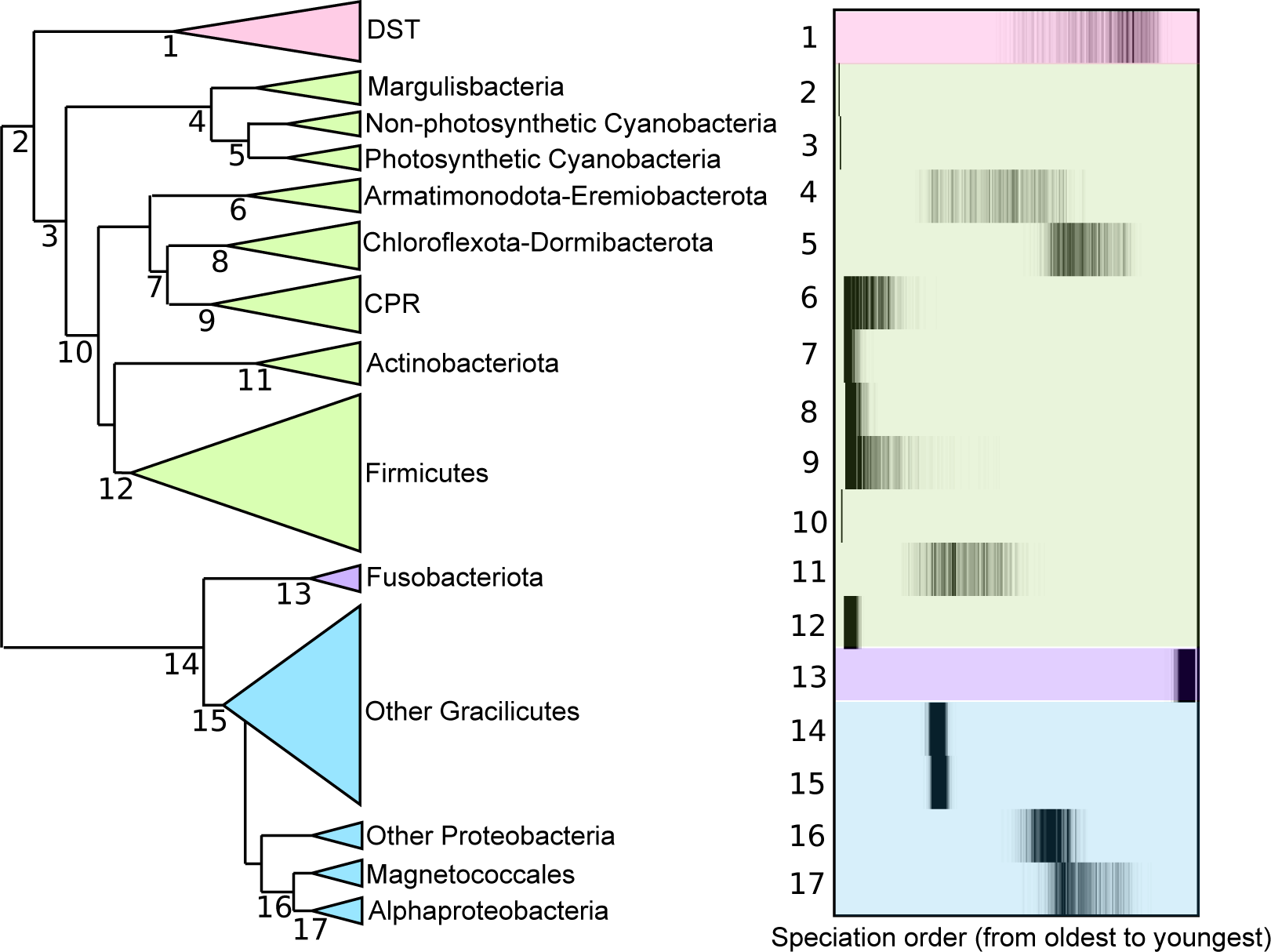
Relative ages of bacterial clades. We used the relative time information provided by directional (donor-to-recipient) patterns of gene transfer to infer the relative ages of bacterial clades. Node numbers in the cladogram (left) correspond to rows in the speciation plot (right). Uncertainties represent the range of sampled time orders that are consistent with high-confidence constraints implied by gene transfers. Following the divergence between Terrabacteria and Gracilicutes, the earliest radiations of extant groups were among Terrabacteria, including CPR and Firmicutes. Note that gene transfers indicate the order of branching events, but provide no information about the absolute time intervals separating events. The geological record provides evidence for oxygenic photosynthesis prior to 3.2Gya^50^, suggesting that divergence 5 - and by extension all earlier divergences - occurred prior to this date. Relative ages for the crown groups of all phyla are provided in Extended Data Figure 4. DST are the Deinococcota, Synergistota, and Thermotogota.

Given the limitations of using a remote archaeal outgroup to establish the root of the bacterial tree, we explored outgroup-free rooting using gene tree-species tree reconciliation^1,35,36,37^. We recently applied this approach to root the archaeal tree^1^, and similar approaches have been applied to investigate the root of eukaryotes^38,39^ and to map and characterise whole genome duplications in plants^40^. The method works by explaining the histories of individual gene families in the context of a shared species tree with a series of speciation, gene origination, duplication, transfer and loss events. Since these histories depend on the position of the root, reconciliation likelihoods can be used to estimate the most likely root, in what can be viewed as a genome-wide extension of the classical approach used to root the tree of life based on ancient gene duplications^4,41^. In addition to leveraging genome-wide data, a further advantage is the ability to extract root signal from both gene duplications and transfers^11,35^. Our method (amalgamated likelihood estimation, ALE) improves on earlier approaches by explicitly accounting for uncertainty in the gene tree topologies and in the events leading to those topologies, while also estimating rates of gene duplication, transfer and loss directly from the data^37^. Simulations suggest that root inferences under ALE are robust to variation in taxon sampling and that the method finds the correct root even under high levels of gene transfer^1,35^, suggesting that the approach is appropriate for the problem at hand (Supplementary Discussion).

We used ALE to test the support for 62 root positions (Extended Data Table 3, Supplementary Table 1) on the unrooted topology by reconciling gene trees for 11,272 homologous gene families from the 265 bacterial genomes. In addition to testing root positions corresponding to published hypotheses, we exhaustively tested all inner nodes of the tree above the phylum level. The ALE analysis rejected all of the roots tested (P < 0.05) except for three adjacent branches, lying between the two major clades of Gracilicutes and Terrabacteria (Figure 1). The position of the phylum Fusobacteriota was difficult to resolve in the tree, and contributed to root uncertainty. The three candidate root branches lead to (i) Terrabacteria+Deinococcus/ Thermotoga/Synergistes; (ii) Gracilicutes; (iii) Fusobacteriota (Figure 1). Consistent with this being the optimal root region, alternative roots were rejected with increasing confidence as distance from the optimal region increased (Figure 1(c), Extended Data Table 3). As taxon sampling and gene family clustering affects phylogenomic inference, we repeated this analysis on a different subsample of representative bacteria and using a different protein clustering approach; these sensitivity analyses (which included a literature-based approach to taxon sampling and COG family clustering) also recovered the root between Gracilicutes and Terrabacteria (Supplementary Methods, Extended Data Figure 3, Supplementary Table 2). A root between Gracilicutes and Terrabacteria was previously reported^42,43^. However, this analysis did not include the CPR, which has been recently suggested^11,16^ to represent the earliest diverging bacterial lineage. Our outgroup-free analysis consistently recovered CPR nested within the Terrabacteria, suggesting that the CPR root is a long branch attraction artifact.

**Figure 3:**
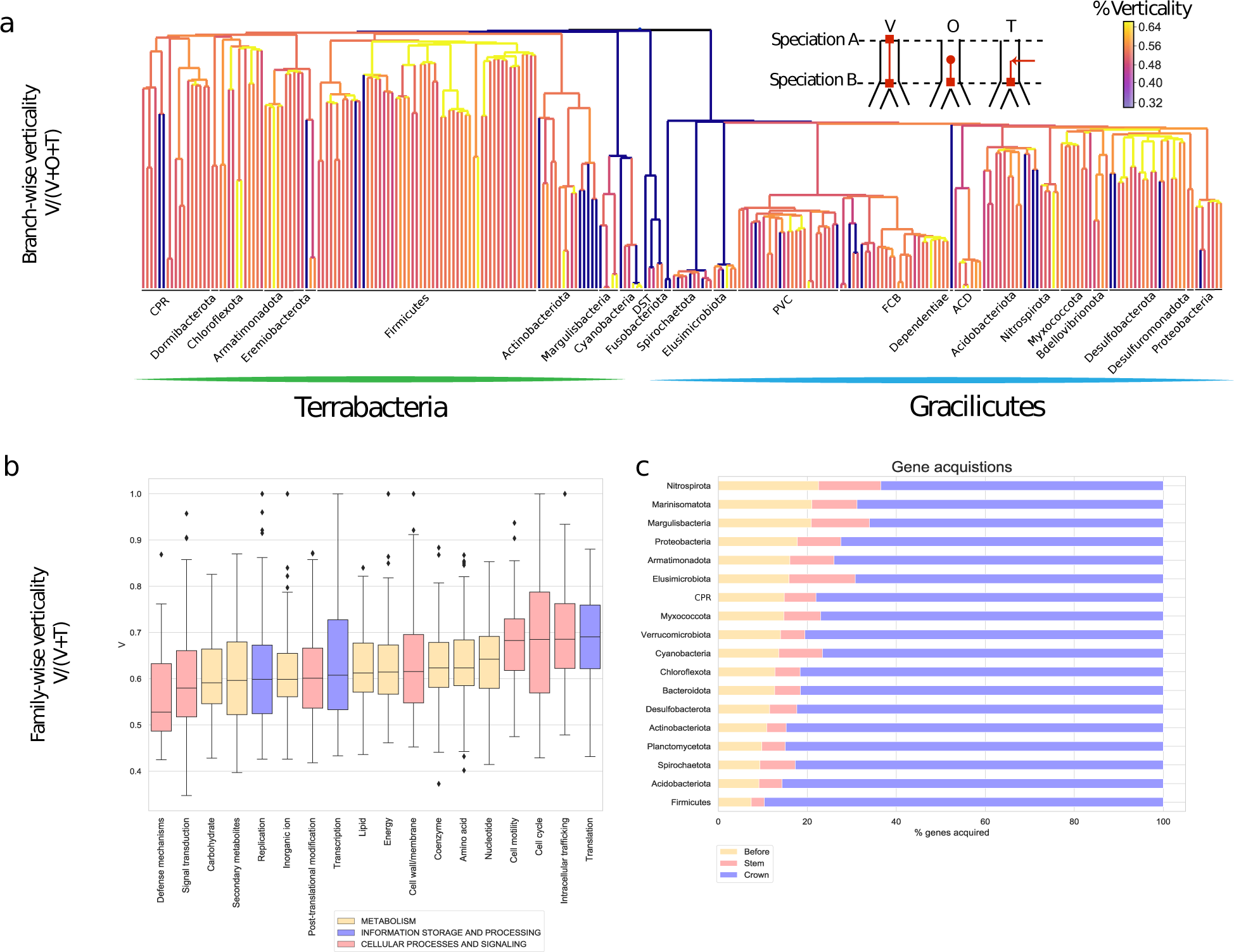
The verticality of bacterial genome evolution. (a) The rooted bacterial species tree (Figure 1) with branches coloured according to verticality: the fraction of genes at the bottom of a branch that descend vertically from the top of that branch (see inset; V = vertical, O = origination, T = transfer into a branch; see Supplementary Methods). Node heights reflect relative time order consistent with highly-supported gene transfers (Figure 2). (b) Verticality by COG functional category: that is, the proportion of gene tree branches that are vertical V/V+T for COG gene families. Genes involved in information processing, particularly translation (J), show the highest verticality (median 0.69), while genes involved in cell defense mechanisms (V, such as genes involved in antibiotic defense and biosynthesis) are most frequently transferred (median verticality 0.53). (c) For a given genome, this combination of vertical and horizontal processes gives rise to a distribution of gene residence times, reflecting the point in evolutionary history at which the gene was most recently acquired. Across all phyla examined, 82% of genes on sampled genomes were most recently acquired after the crown group radiation of that phylum. FCB are the Fibrobacterota, Chlorobiota, Bacteroidota, and related lineages; PVC are the Planctomycetota, Verrucomicrobiota, Chlamydiota, and related lineages; DST are the Deinococcota, Synergistota, and Thermotogota; ACD are Aquificota, Campylobacterota, and Deferribacterota.

The rooted tree (Figure 1) indicates that the CPR are a derived group that branches sister to Chloroflexota-Dormibacterota, and likely evolved from free-living ancestors. The absence of an electron transport chain in many CPR^19^ is therefore likely to result from secondary loss in the CPR common ancestor.

Despite being a derived lineage within Terrabacteria, patterns of gene transfer suggest that the radiation of CPR was one of the earliest events during the diversification of Bacteria (Figure 2). Transfers contain information about the relative timing of divergences because donors must be at least as old as their recipients^44,45^. Since inferred transfer events are uncertain, we used only high-confidence relative age constraints recovered in at least 95/100 bootstrap replicates (see Supplementary Methods) to establish the relative ages of bacterial clades (Figure 2, Extended Data Figure 4). These analyses suggest that several groups within Terrabacteria are older than the entire Gracilicutes radiation, including the crown groups of the CPR (97.4% of sampled time orders) and Firmicutes (100% of sampled time orders). By contrast, the emergence of the Alphaproteobacteria and the photosynthetic Cyanobacteria were relatively late events during bacterial evolution: the divergence between Alphaproteobacteria and *Magnetococcales* was the 153rd of 264 internal divergences (median rank), while the divergence of photosynthetic Cyanobacteria from their closest relatives had a median rank of 172nd (Figure 2). These divergences confirm that the mitochondrial and plastid endosymbioses, and therefore the origin of eukaryotic cells, occurred during the later stages of bacterial diversification^46–50^.

**Figure 4:**
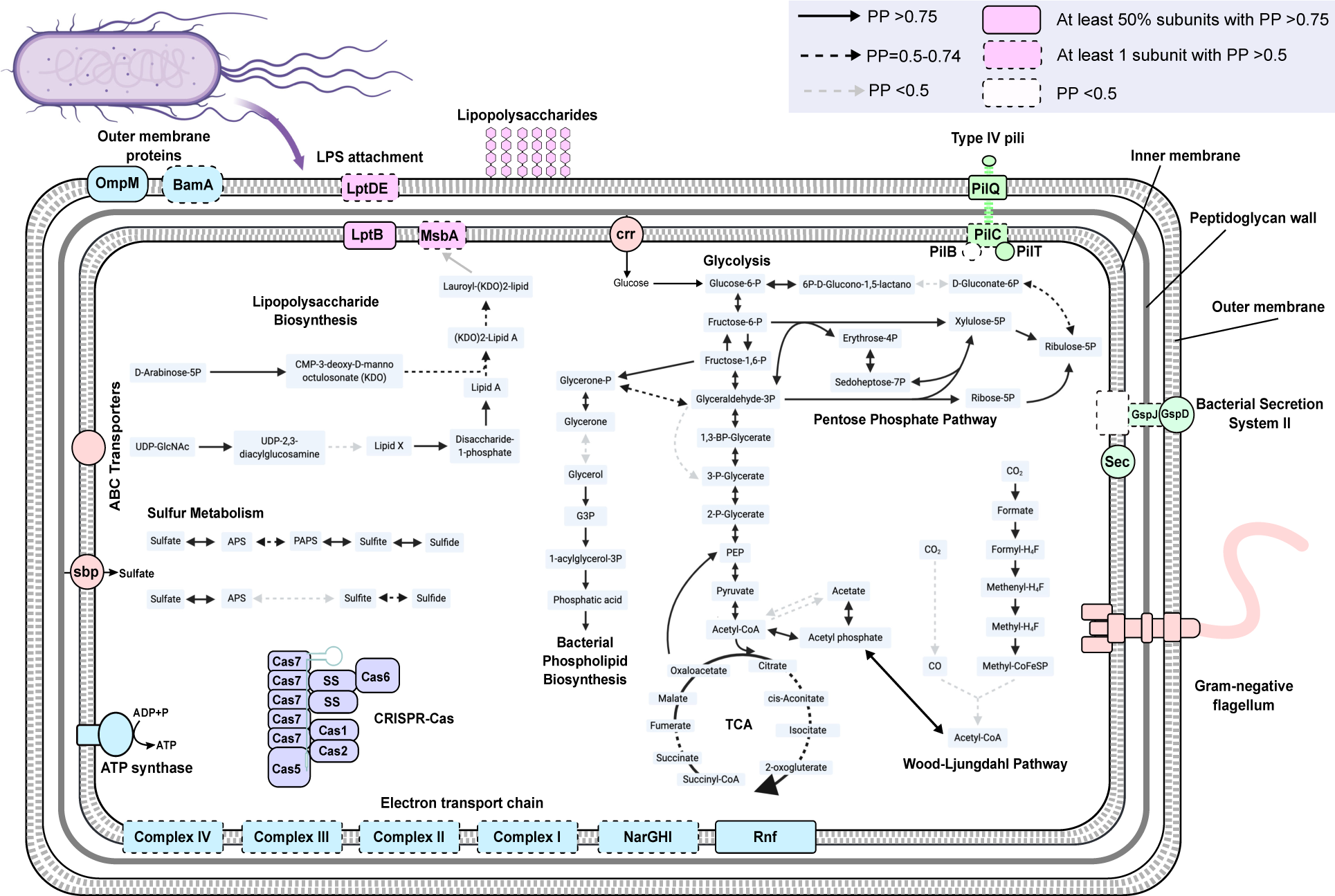
Ancestral reconstruction of the last bacterial common ancestor (LBCA). The reconstruction is based on genes that could be mapped to at least one branch within the root region with PP >0.5 (see Supplementary Discussion and Extended Data Figure 7, Extended Data Figure 8). The presence of a gene within a pathway is indicated as shown in the key. Our analyses suggest that LBCA was a rod-shaped, motile, flagellated double-membraned cell. We recover strong support for central carbon pathways, including glycolysis, the tricarboxylic acid cycle (TCA) and the pentose phosphate pathway. We did not find unequivocal evidence for the presence of a carbon fixation pathway, although we found moderate support for components of both the Wood-Ljungdahl pathway and the reverse TCA cycle. Though not depicted here, our analyses suggest that the machinery for transcription, translation, tRNA and amino acid biosynthesis, homologous recombination, nucleotide excision and repair, and quorum sensing was also present in LBCA (see Supplementary Tables 4 and 5).

## Is bacterial evolution treelike?

How much of bacterial evolution can be explained by the concept of a rooted species tree? Horizontal gene transfer (HGT) is frequent in prokaryotes, and published analyses indicate that most or all prokaryotic gene families have experienced HGT during their history^1,51^. This implies that there is no single tree that fully describes the evolution of all bacterial genes or genomes^52,53^. Extensive HGT is existentially challenging for concatenation, because it greatly curtails the number of genes that evolve on a single underlying tree^54^. Phylogenetic networks^53,55^ were the first methods to explicitly acknowledge non-vertical evolution, but can be difficult to interpret biologically. Gene tree-species tree reconciliation integrates tree and network-based approaches by modelling both the horizontal components of genome evolution (a fully reticulated network allowing all possible transfers) and the vertical trace (a common rooted species tree). This framework enables us to quantify the contributions of vertical and horizontal processes to bacterial evolutionary history.

Our analyses (Figure 3) reveal that most bacterial gene families present in at least two species (9678/10518 MCL families, 92%) have undergone at least one gene transfer during their evolution; only very small families have escaped transfer entirely (Extended Data Figure 5). Note that our inference of HGT events is certain to be an underestimate, because our broad but sparse taxon sampling does not allow us to detect transfers or recombination within strains. Consistent with previous analyses^1,56^, transfer rates vary across gene functional categories, with genes functioning in defense mechanisms (such as antibiotic biosynthesis) and the production of secondary metabolites being the most frequently transferred, and those involved in translation and the cell cycle the least (Figure 4(b)). Despite this accumulation of HGT, most gene families evolve vertically the majority of the time, in that 68% of transmission events (mean, MCL gene families) are vertical along the species tree.

Mapping the branches of the gene trees onto the species tree demonstrates that the optimal tree provides an apt summary of much of bacterial evolutionary history, even for the deepest branches of the tree^57–59^. From the gene’s eye view, gene families evolve neither entirely vertically nor horizontally: core genes are occasionally transferred, and even frequently exchanged genes contribute useful vertical signal; for example, the median number of genes that evolve vertically on a branch of the species tree is 998.92 (Supplementary Table 3), far greater than the number of genes that have been concatenated at the level of all Bacteria. From the perspective of the genome, constituent genes have different ages, corresponding to the time at which they originated or were most recently acquired by gene transfer, within the resolution of our taxonomic sampling. This analysis indicates that, on average, 82% of the genes on all genomes from adequately represented phyla (5 or more genomes) were most recently acquired after the diversification of that phylum, though all genomes retain a smaller proportion (10.3-26.7%) of genes that have descended vertically from the stem lineage of their phylum or even earlier (Figure 3(c)).

## Ancestral proteome of the Last Bacterial Common Ancestor (LBCA)

Reconciliation analysis not only allows us to infer the acquisition of genes across the tree, but also to estimate the metabolic potential of the LBCA. To do so, we built a second smaller set of gene families from COG annotations, which are better suited for functional annotation, and reconciled their gene trees with the species tree (see Supplementary Methods). Protein family annotations (COG and KO) and root presence posterior probabilities (PPs) for all 3723 gene families under all three roots are provided in Supplementary Table 4; in the following reconstruction, we indicate when gene content inferences differ between roots. PPs for genes directly relevant to our reconstruction are provided in Supplementary Table 5. Based on the root placement and estimated rates of gene family extinction^1^, we predict that LBCA encoded 1292.6-2142.9 COG family members, the majority of which (median estimates 65-69.5%; 95% CI 57-82%) survived to be sampled in at least one present day genome. Based on the relationship between COG family members and genome size for extant Bacteria (Pearson’s r = 0.96, *P =* 8 × 10^−153^), we estimate the genome size of LBCA to be 2.69Mb +/-0.4Mb (standard error) for root 1 (Fusobacteriota with Terrabacteria; Figure 1(b)); 2.59Mb +/-0.41Mb for root 2 (Fusobacteriota with Gracilicutes), and 1.6 +/-0.5Mb for root 3 (Fusobacteriota basal). Under all three roots, the trend in genome size evolution from LBCA to modern taxa is an ongoing moderate increase through time in estimated COG family complements and genome sizes. Genome reduction of 0.47-0.56Mb on the CPR stem lineage after divergence from their common ancestor with Chloroflexota is the most significant departure from this trend (Extended Data Figure 6). COG families lost on the CPR stem include components of the electron transport chain, carbon metabolism, flagellar biosynthesis and motor switch proteins, amino acid biosynthesis, the Clp protease subunit ClpX and RNA polymerase sigma factor-54, in agreement with previous findings^18^ (Supplementary Table 6).

As might be expected, this ancestral gene set includes most of the components of the modern bacterial transcription, translation and DNA replication systems. It also includes an FtsZ-based cell division machinery and pathways for signal transduction, membrane transport and secretion (Figure 4, Supplementary Discussion). Further, we identified proteins involved in bacterial phospholipid biosynthesis, suggesting that LBCA had bacterial-type ester-lipid membranes (Figure 4). We also identified most of the proteins required to synthesize appendages such as flagella and pili as well as to enable quorum sensing, suggesting that LBCA was motile; which is in agreement with the previous suggestion that flagella were present in LBCA^60^. Since bacterial genes are typically maintained by strong positive selection^61^, these findings imply that LBCA lived in an environment in which dispersal, chemotaxis and surface attachment were advantageous.

Moderate support for the presence of *mreB* (0.9/0.88/0.73, root branches 1-3 as depicted in Figure 1(b)), *mreC* (0.82/0.79/0.57) and *mreD* (0.86/0.83/0.63) at the root suggests that LBCA possessed rod-shaped cells. We also obtained high root posterior probabilities for proteins mediating outer cell envelope biosynthesis including for lipopolysaccharides (LPS), from which we infer that LBCA possessed a double membrane with an LPS layer (Supplementary Discussion). Consistent with this inference, we obtained high posterior probabilities for the flagellar subunits FlgH, FlgI and FgA in LBCA, which anchor flagella in diderm membranes^62^, and for the Type IV pilus subunit PilQ, which among extant bacteria is specific to diderms^62,63^. Altogether, this is consistent with hypotheses^9^ in which LBCA was a diderm^62,63^, and argues against scenarios in which the Gram-negative double membrane originated by endosymbiosis between monoderms (single-membraned bacteria^10^) or via the arrest of sporulation^64^ in a spore-forming monoderm ancestor. Subsequently, diderm-to-monoderm transitions may have occurred on multiple occasions within Bacteria^62,63^.

We recovered components of several core pathways for carbohydrate metabolism with high posterior support, including glycolysis, the tricarboxylic acid (TCA) cycle, and the pentose phosphate pathway (Figure 4, Extended Data Figure 7, Extended Data Figure 8, Supplementary Table 5, Supplementary Discussion). Modern bacteria fix carbon using several different pathways, including the Calvin cycle, the 3-hydroxypropionate bicycle and variations thereof as well as the Wood-Ljungdahl Pathway (WLP) and the reverse TCA cycle, the latter two of which have been suggested to have emerged early in the history of life^43,65–69^. Of these, we identified several enzymes of the TCA cycle, although the directionality of the enzymes is difficult to assess^70^ (Figure 4, Extended Data Figure 7, Extended Data Figure 8, Supplementary Discussion). Furthermore, we identified several enzymes of the methyl-branch of the WLP, for acetate biosynthesis as well as components of a putative RNF complex (Figure 4, Extended Data Figure 7, Extended Data Figure 8), which together may indicate that LBCA was capable of acetogenic growth^71^ (Supplementary Discussion). However, the key enzyme of the WLP, the Carbon monoxide dehydrogenase/acetyl-CoA synthase complex^43^, had only moderate root support (PP=0.5-0.75 for two subunits) and low support (PP <0.5) for other subunits. Thus, while our analyses provide strong support for the antiquity of components of the WLP, acetogenesis, the TCA cycle and several other core metabolic pathways, they do not confidently establish the combination of pathways employed by LBCA as distinct from other organisms present at the same time (Supplementary Discussion).

Finally, our reconstruction also indicated high posterior support for several elements of an adaptive immunity CRISPR-Cas system^72,73^, including the universally conserved Cas endonuclease, Cas1 (PP=0.96/0.93/0.89), which is essential for spacer acquisition and insertion into CRISPR cassettes^74,75^. Interestingly, highly supported CRISPR components in LBCA belong primarily to Class 1 systems, specifically Type I and Type III, which exhibit greater modular diversity than their Class 2 counterparts (Supplementary Table 4, Supplemental Information)^73^. We recovered a near complete prototypical Type III CRISPR system (Supplementary Table 4, Supplementary Information), providing strong support for its presence in LBCA. Among other roles, CRISPR systems are crucial in antiviral defense and activate in response to viral exposure^76^; therefore these findings are consistent with hypotheses suggesting that LBCA already co-evolved with parasitic replicators such as bacteriophages^77 78^.

## Conclusions

Our analyses suggest that, despite extensive horizontal gene transfer, a phylogenetic tree is an apt representation of bacterial evolution in the sense that most bacterial gene families evolve vertically most of the time. In contrast to recent outgroup-rooted analyses, we found no support for a root on the CPR branch; instead, our analysis suggests that this lineage evolved from a common ancestor with Chloroflexota by reductive evolution, and that these and other divergences within the Terrabacteria were some of the earliest events in bacterial diversification. We place the last bacterial common ancestor between two major clades, Terrabacteria and Gracilicutes, although we could not resolve the position of Fusobacteriota in relation to those major radiations. Fusobacteriota currently comprise anaerobic free-living, pathogenic and commensal diderm bacteria^79^, and a clear direction for future work will be to place them on the rooted bacterial tree, particularly if more basal members of this lineage come to light. Ancestral gene content reconstruction on the rooted tree suggests that LBCA was a free-living, fully fledged diderm with LPS, a multimeric flagellum, and a type III CRISPR-Cas system presumably for defence against viruses. Further progress on these questions will require developments on at least two fronts. Environmental genomics has radically expanded our sampling of prokaryotic diversity, but despite enormous progress we have sampled only ∼30,000 of the estimated 2-4 million^80^ prokaryotic species in the biosphere: there is much more diversity out there to discover. At the same time, current phylogenomic methods neither capture the full complexity of evolution nor scale computationally to analyze the genomes discovered so far. As more diversity comes to light, improved reconciliation methods will continue to aid our understanding of LBCA and subsequent major evolutionary transitions across the tree of life.

## Methods

### Taxon sampling and unrooted species tree inference

To obtain a representative taxon sampling from across known bacterial diversity, we sampled taxa according to the classification provided by the Genome Taxonomy Database (GTDB), obtaining 265 genomes (see Supplementary Methods). We used OMA 2.1.1^24^ to identify candidate single-copy bacterial orthologues, and retained those with at least 75% of all species represented in each family. Sequences were aligned in Mafft using the -auto option, and trimmed in BMGE 1.12^81^ using the BLOSUM30 model. Initial trees were inferred for each candidate marker gene under the LG+G+F model in IQ_TREE 1.6.10. The trees were manually inspected, and we selected orthologues where the monophyly of 14 pre-defined major lineages was not violated with bootstrap support >70%, resulting in 63 final orthologues (see Supplementary Methods). Concatenation of this marker set resulted in an alignment of 18,234 amino acids. We inferred an unrooted phylogeny from this concatenate under the LG+C60+R8+F model, which was chosen as the best-fitting model by the BIC criterion in IQ-TREE^82^. We additionally removed the most compositionally heterogeneous sites from the sequence alignment using Alignment Pruner^83^ (https://github.com/novigit/davinciCode/blob/master/perl) (20%, 40%, 60% and 80% respectively) and inferred trees using the same procedure described above in order to compare the resulting topologies (see Supplementary Methods).

### Outgroup rooting

To root the bacterial tree using an archaeal outgroup, we used a representative sampling of 148 archaeal genomes and inferred the ML tree in IQ-TREE under the best-fitting LG+C60+R8+F model. The concatenated alignment included a subset of 30/ out of the 63 bacterial orthologues that were shared between bacteria and archaea, as determined by HMM searches and manual inspection of single gene trees. We performed approximately-unbiased (AU) to determine whether a range of published alternative rooting hypotheses (Table 1) could be rejected, given the model and data (AU p-value > 0.05).

### Gene family clustering and ALE analysis

To infer homologous gene families for ALE inference (ALE v0.4), we performed an all vs all similarity search using Diamond v0.9.25^84^ with an E-value threshold of <10^−7^, and clustering using MCL with an inflation parameter of 1.2. This resulted in 186,827 gene families and a total of 11,765 families with 4 or more sequences. Sequence alignment and trimming was performed as described above. After site filtering, 260 alignments that contained no high-quality columns were discarded as were all alignments with less than 30 columns, resulting in 11,272 alignments. We subsequently filtered out sequences containing more than 80% of gaps to produce the final set of alignments. The trees were inferred using IQ-TREE under the best-fitting model (as determined by BIC), and we generated 10000 rapid bootstrap replicates per gene. Conditional clade probabilities^85^ (CCPs) were computed using ALEobserve and the resulting ALE files were reconciled with the species tree. Loss rates were corrected by genome completeness, estimated using the values of CheckM^86^ (https://data.ace.uq.edu.au/public/gtdb/data/releases/release89/89.0/bac120_metadata_r89.tsv) We estimated reconciliation likelihoods for 62 candidate roots on the unrooted species tree (see Supplementary Information).

#### Inference of relative divergence times of bacterial clades

We parsed the transfers inferred using ALEml_undated (from the output uT files; they can be found in the Extended Data, ReconciliationsMCL.zip) and discarded those with posterior probability < 0.05. We used bootstrapping to estimate constraint support in the following way: for each of the three candidate species trees, we sampled the gene families 100 times with replacement and, for each replicate, converted detected transfers to constraints and performed a MaxTiC analysis^44,87^. A total of 8743, 8629, 9079 constraints were recovered in at least 95/100 replicates for the 3 possible roots respectively, and we used this subset of highly supported constraints in our final analysis. We generated 1000 time orders compatible with those constraints for every root using the script mc_explorer.py (Extended Data Files). We then ranked all interior nodes on the tree, with the root node having rank 0 and the most recent speciation node having rank 263.

### Ancestral gene content and metabolic reconstruction

We clustered proteins into families based on their COG annotations, resulting in 3723 COG families with at least 4 sequences (see Supplementary Information). We estimated the posterior probability of presence at the root node for each gene family under the gene tree-species tree reconciliation analysis using root origination rates estimated independently for each of the 23 COG functional categories. Initial gene content and metabolic inferences at a particular node were based on gene families with a posterior presence probability (PP) of >0.95 at that node, with further manual inspection of PPs for incomplete pathways (see Supplementary Information for more details). We also considered the PPs of the components of the same metabolic pathway or functional module for genes with moderate root posterior probabilities (0.5 < PP < 0.95) to evaluate gene pathway and root presence. There are many pathways for which most or all components have moderate to high posterior support (PP >0.5), and these pathways are likely to have been present in LBCA. See Supplemental Discussion and Data for all PPs and further discussion of gene- and pathway-level support.

### Protein and protein family functional annotation

Protein sequences from all genomes used for phylogenetic analyses in this study were annotated using a variety of databases. Functional annotations were obtained using hmmsearch v3.1b2 (settings: -E 1e-5) ^88^ against KOs from the KEGG Automatic Annotation Server (KAAS; downloaded April 2019) ^89^. Additionally, all proteins were scanned for protein domains using InterProScan (v5.31-70.0; settings: --iprlookup --goterms)^90^. Multiple hits corresponding to the i n d i v i d u a l d o m a i n s o f a p r o t e i n a r e r e p o r t e d u s i n g a c u s t o m s c r i p t (parse_IPRdomains_vs2_GO_2.py). For the functional annotation of the 4256 COG families investigated in our ancestral reconstructions, we assigned KOs using a majority rule: i.e. we assigned the KO, that was reported in > 50% of the sequences comprising each of the COG families yielding a COG-to-KO mapping file. Subsequently, we mapped COG descriptions, COG Process/Class, Category description, kegg id, kegg description, and kegg pathway to the COG-to-KO mapping file. COG descriptions were collected from the root annotations (1_annotations.tsv) downloaded at EggNOG (v5.0.0)^91^. COG functional category and Process/ Class descriptions were derived from EggNOG (v4.0) ^92,93^. KO pathways were manually curated based on an in-house KO-to-pathway mapping file, and were subsequently mapped to the respective KO. The scripts for annotation and mapping are included in the Data Supplement.

### Metabolic comparisons

Results from the PP analysis were used as the framework for metabolic comparisons and reconstruction of the proteome of LBCA. First, the occurrence of an individual COG family across each taxon was counted in R (v3.6.3) (Supplementary Table 4). This binary presence/ absence matrix was combined with the PP values for Nodes corresponding to the CPR, Chloroflexota+CPR, Chloroflexota, Terrabacteria, DST+Terrabacteria, Gracilicutes-Spirochaetota, Gracilicutes+Spirochaetota, Root 1, Root 2, and Root 3, filtered with a cutoff of PP>0.50. The combined count table was summarized using the ddply function of the plyr package (v1.8.4), which was used to summarize the counts across each phylogenetic cluster, node, and root. Data is visualized in a heatmap generated using the ggplot function with geom_tile and facet_grid of the ggplot2 package (v3.2.0). Heat map categories for pathways were scaled based on the number of COG families, results were plotted using the grid.draw function of the grid package (v3.6.3). Heatmaps were manually merged with a representative tree in Adobe Illustrator (v22.0.1).

### Quantifying vertical and horizontal signals in bacterial genome evolution

In the context of our analyses, “verticality” is the proportion of inferred evolutionary events that reflect vertical descent, estimated using gene tree-species tree reconciliation. We considered two kinds of verticality: branch-wise verticality, the proportion of vertical evolutionary events on a branch in the species tree; and family-wise verticality, the proportion of vertical events during the evolution of a specific gene family. We defined branch-wise verticality as V/(V+O+T), where V is the inferred number of vertical transmissions of a gene from the ancestral to descendant ends of the branch; O is the number of new gene originations on the branch; and T is the number of gene transfers into the branch. We defined family-wise verticality as V/(V+T), where V and T refer to inferred numbers of events within the history of a gene family (Supplementary Table 7). The numbers reported in the manuscript have been averaged over the reconciliations obtained using the three possible roots.

## Data and code availability

All data and code, including implementations of new methods, are provided online at DOI 10.6084/m9.figshare.12651074.

## Author contributions

The project was conceived by TAW, GJSz, PH, AS, GAC and AAD. GAC, AAD, TAW and GSz performed phylogenomic analyses. GJSz developed new analytical methods. TM, GAC, AAD and AS performed metabolic annotations and reconstructions. All authors contributed to interpretation and writing.

## Acknowledgements

GAC is supported by a Royal Society Research Grant to TAW. TAW is supported by a Royal Society University Research Fellowship and NERC grant NE/P00251X/1. GJSz received funding from the European Research Council under the European Union’s Horizon 2020 research and innovation program under Grant Agreement 714774 and Grant GINOP-2.3.2.-15-2016-00057. AS was supported by the Swedish Research Council (VR starting grant 2016-03559 to AS) and the NWO-I foundation of the Netherlands Organisation for Scientific Research (WISE fellowship to AS). AAD and PH were supported by an Australian Research Council Laureate Fellowship (grant no. FL150100038).

