## Supplementary Information (text and figures) for "A rooted phylogeny resolves early bacterial evolution"

### A rooted phylogeny resolves early bacterial evolution (Supplementary Information)

All of the files referred to below are provided in the data supplement (extended data files) to our paper, available in the FigShare repository at DOI 10.6084/m9.figshare.12651074.

#### Supplementary Data

##### Extended Data Figures

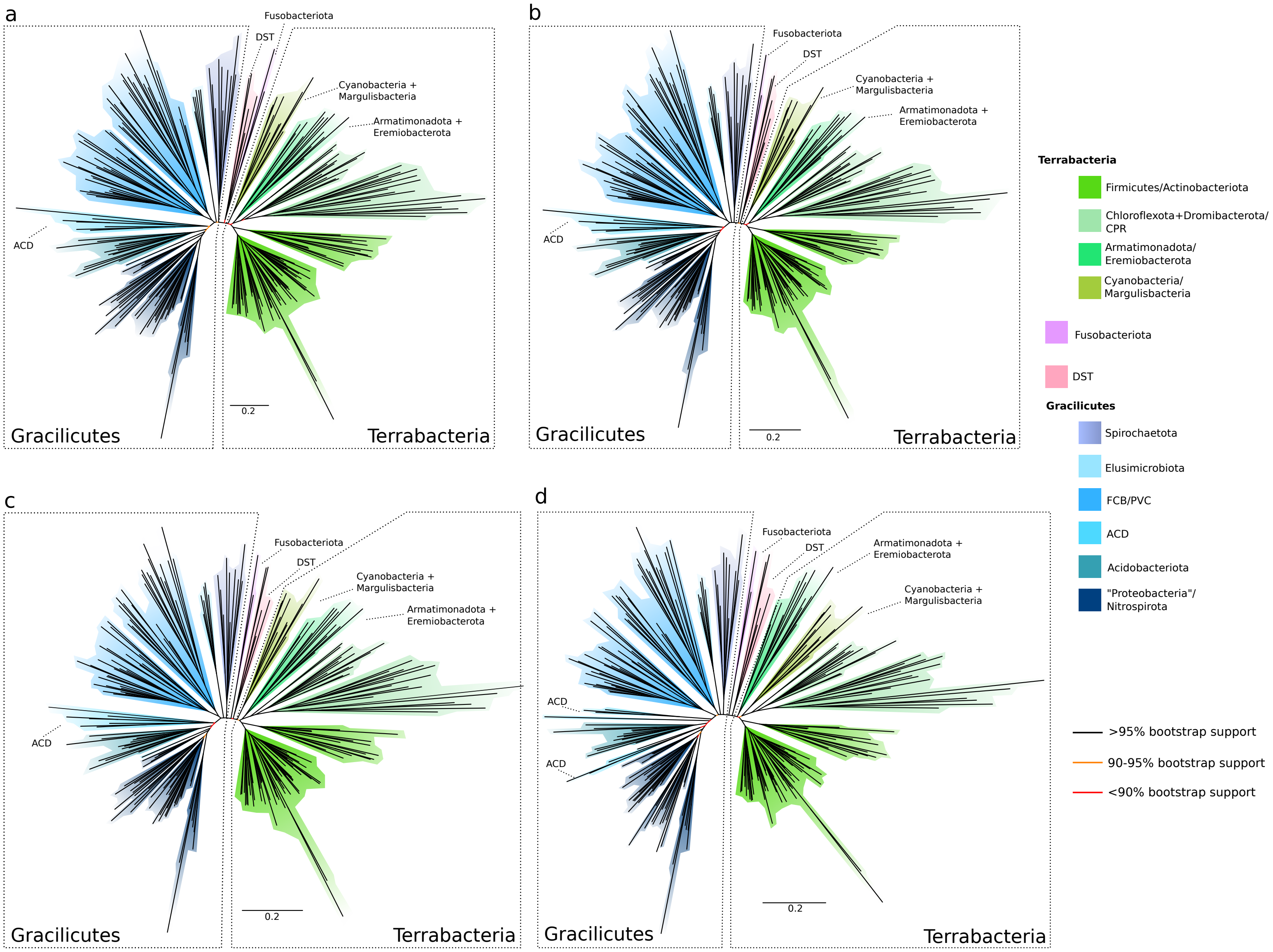

a
Fusobacteriota
DST
Cyanobacteria +
Margulisbacteria
Armatimonadota +
Eremiobacterota
ACD
0.2
Gracilicutes
Terrabacteria
b
Fusobacteriota
DST
Cyanobacteria +
Margulisbacteria
Armatimonadota +
Eremiobacterota
ACD
0.2
Gracilicutes
Terrabacteria
c
Fusobacteriota
DST
Cyanobacteria +
Margulisbacteria
Armatimonadota +
Eremiobacterota
ACD
0.2
Gracilicutes
Terrabacteria
d
Fusobacteriota
DST
Armatimonadota +
Eremiobacterota
Cyanobacteria +
Margulisbacteria
ACD
ACD
0.2
Gracilicutes
Terrabacteria
Terrabacteria
Firmicutes/Actinobacteriota
Chloroflexota+Dromibacterota/
CPR
Armatimonadota/
Eremiobacterota
Cyanobacteria/
Margulisbacteria
Fusobacteriota
DST
Gracilicutes
Spirochaetota
Elusimicrobiota
FCB/PVC
ACD
Acidobacteriota
"Proteobacteria"/
Nitrospirota
>95% bootstrap support
90-95% bootstrap support
<90% bootstrap support

**Extended Data Figure 1: Maximum likelihood unrooted bacterial phylogeny under the best-fitting substitution model (LG+C60+R8+F) following removal of the 20%-80% most compositionally heterogeneous sites.** Sites were identified and removed using Alignment Pruner. (a) 20% most compositionally heterogeneous removed, with 14580/18234 sites remaining following site stripping; (b) 40% most compositionally heterogeneous removed, with 10941/18234 sites remaining following site stripping; (c) 60% most compositionally heterogeneous removed, with 7294/18234 sites remaining following site stripping; (d) 80% most compositionally heterogeneous removed, with 3647/18234 sites remaining following site stripping; Branch supports are ultrafast bootstraps, branch lengths are proportional to the expected number of substitutions per site.

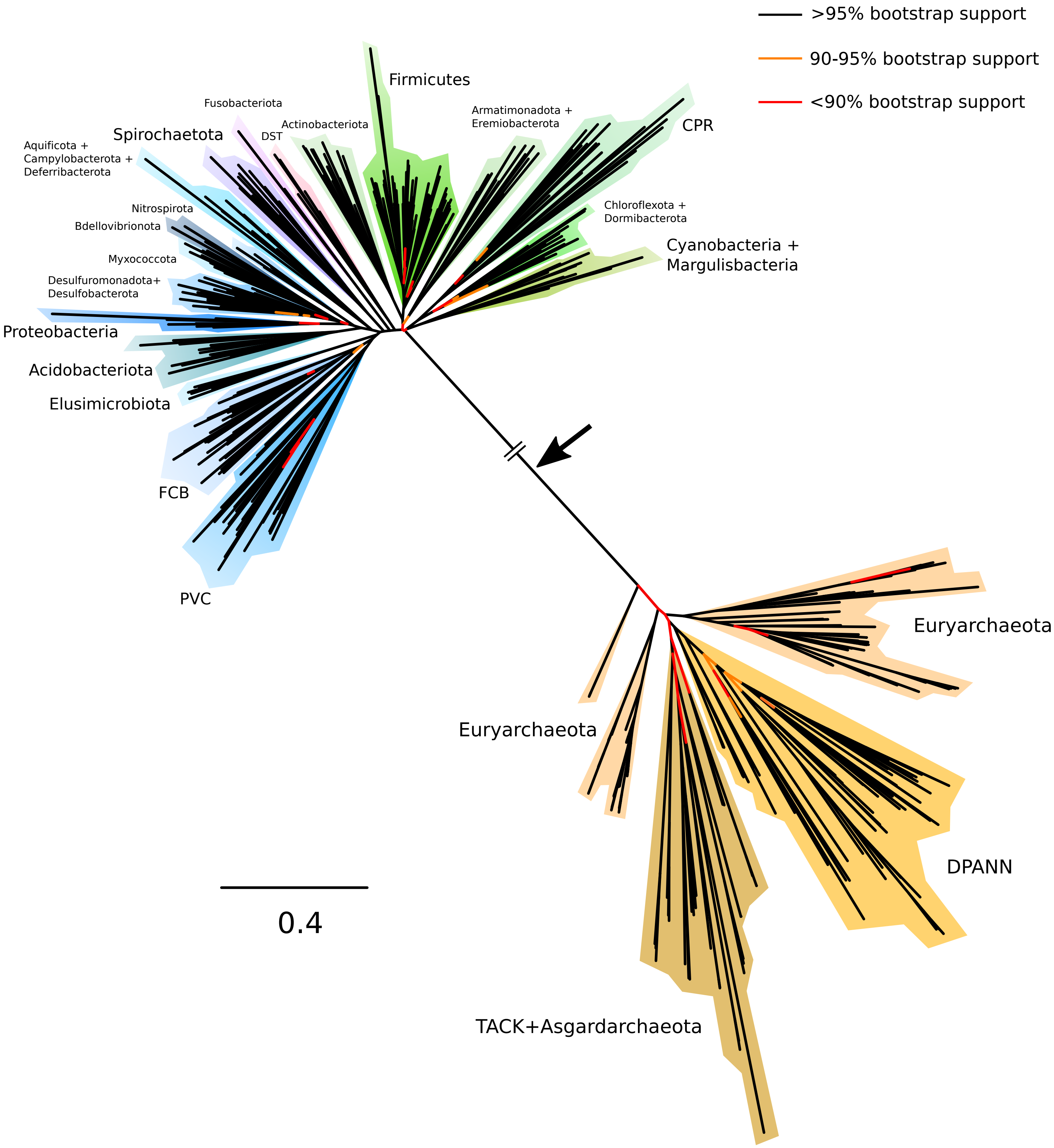

>95% bootstrap support
90-95% bootstrap support
<90% bootstrap support
Firmicutes
Fusobacteriota
Actinobacteriota
DST
Spirochaetota
Aquificota +
Campylobacterota +
Deferribacterota
Armatimonadota +
Eremiobacterota
CPR
Chloroflexota +
Dormibacterota
Cyanobacteria +
Margulisbacteria
Nitrospirota
Bdellovibrionota
Myxococcota
Desulfuromonadota+
Desulfobacterota
Proteobacteria
Acidobacteriota
Elusimicrobiota
FCB
PVC
Euryarchaeota
Euryarchaeota
DPANN
TACK+Asgardarchaeota
0.4

**Extended Data Figure 2: Maximum likelihood outgroup-rooted bacterial phylogeny.** The maximum likelihood phylogeny obtained under the best-fitting LG+C60+R8+F model on a concatenation of 30 marker genes shared between Bacteria and Archaea. The bacterial root (marked by a black arrow) separates CPR, Cyanobacteria+Margulisbacteria, and Chloroflexota+Dormibacterota from the rest of the bacterial tree, but this position has poor bootstrap support and a range of alternative hypotheses could not be rejected statistically; note also that a basal position for DPANN within Archaea[1,2] could not be rejected using an Approximately Unbiased (AU) test (**Extended Data Table 2**). FCB are the Fibrobacterota, Chlorobiota, Bacteroides, and related lineages; PVC are the Planctomycetes, Verrucomicrobia, Chlamydiae, and related lineages; DST are the Deinococcota, Synergistota, and Thermatogota; ACD are Aquificota, Campylobacterota, and Deferribacterota; FA are Firmicutes and Actinobacteria. Branch supports are ultrafast bootstraps, as indicated by the colour key. Branch lengths are proportional to the expected number of substitutions per site.

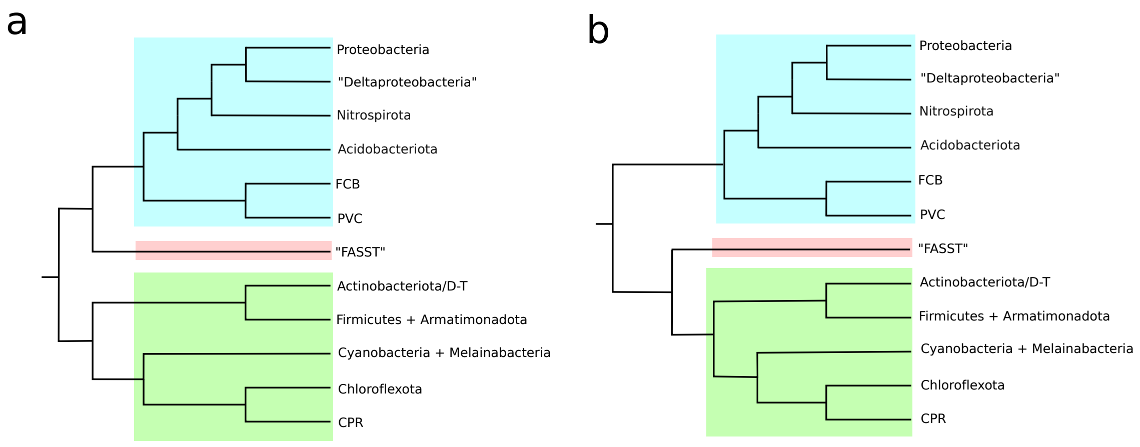


**Extended Data Figure 3: Two rooted topologies from the GTDB-independent sensitivity analysis that could not be rejected by the AU test, from ALE analysis incorporating genome completeness.** AU p-values are 0.973 for tree (a) and 0.064 for tree (b). Both trees are in agreement with each other and with the focal analysis in placing the root between Terrabacteria and Gracilicutes, but disagree in the placement of the "FASST" taxa comprising Fusobacteriota, Aquificota, Synergistota, Spirochaetota and Thermotogota. D-T stands for Deinococcus-Thermus; "Deltaproteobacteria" is Desulfuromonadota, Desulfobacterota, Bdellovibrionota, and Myxococcota.

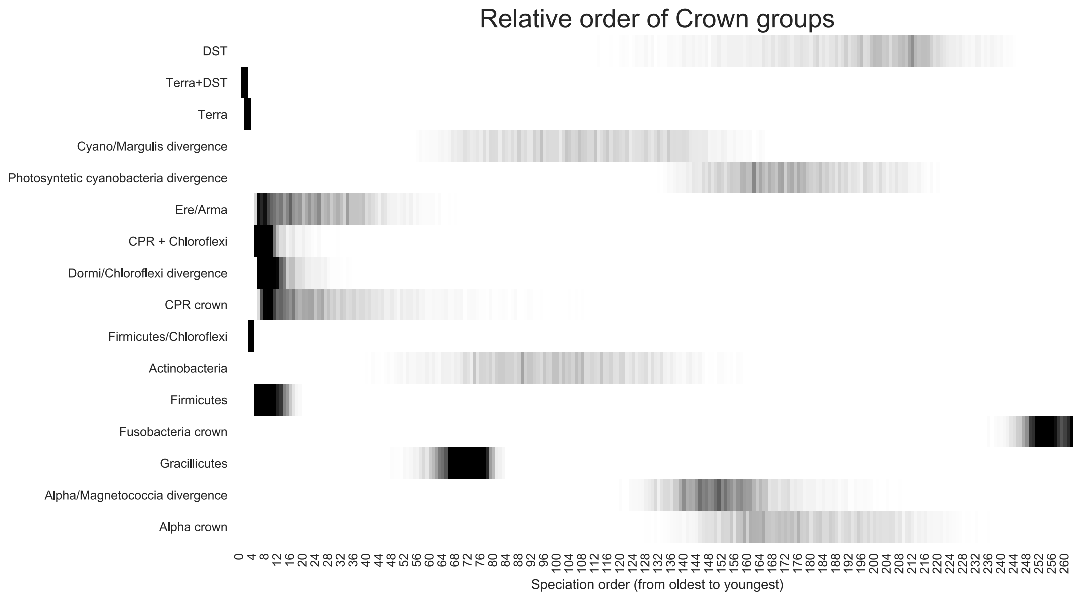


**Extended Data Figure 4: Relative ages for the crown groups of bacterial phyla.** The relative ages were inferred by generating random time orders that were fully compatible with all highly supported constraints (see Supplementary Methods). Speciations are ordered from oldest (the root) to most recent. When interpreting this plot, it is important to note that time orders are relative, and the analysis does not contain any information about the absolute amount of geological time that elapsed between any two speciation events. Only phyla represented by at least two genomes are included in the plot.

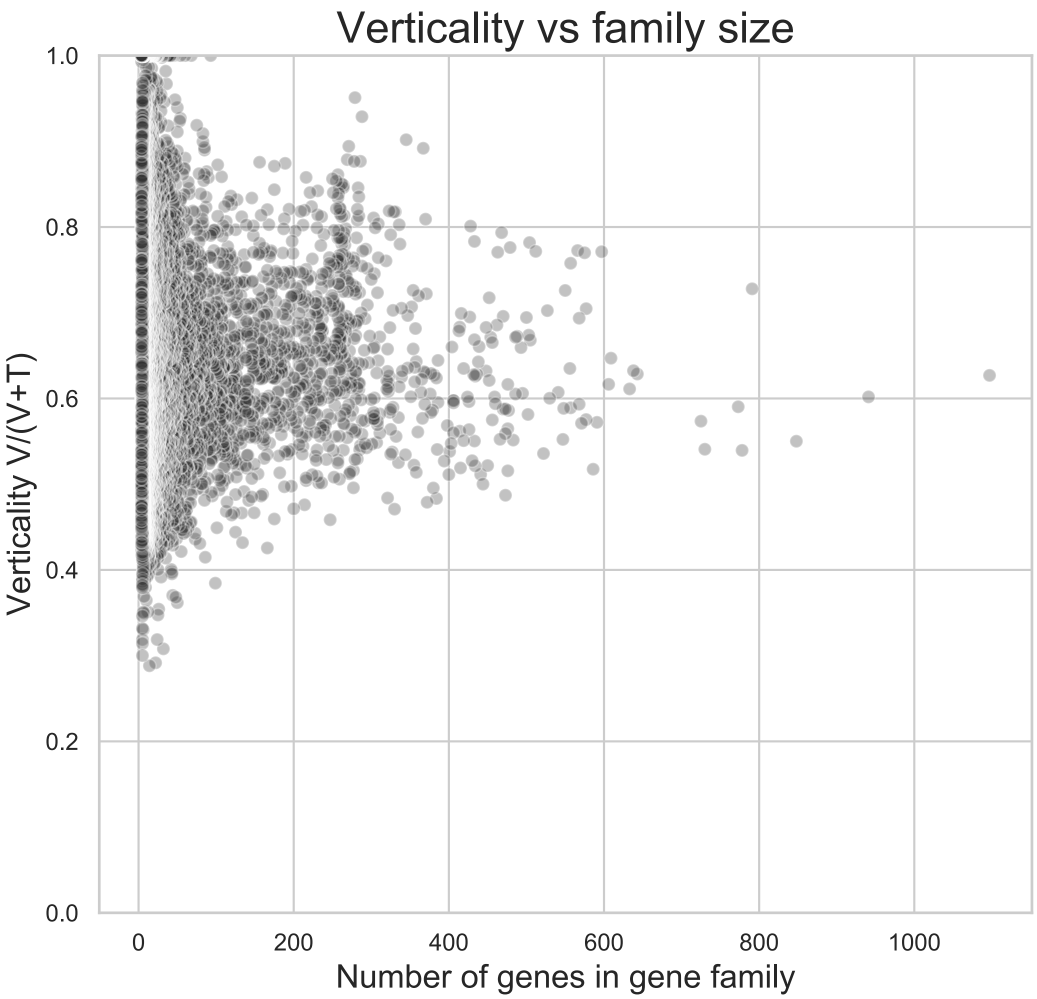


**Extended Data Figure 5: The relationship between verticality and gene family size.** Most gene families have experienced many transfers. Verticality varies with gene functional class, but families with very low transfer rates are small; these might represent young families that have not yet had enough time to experience gene transfer.

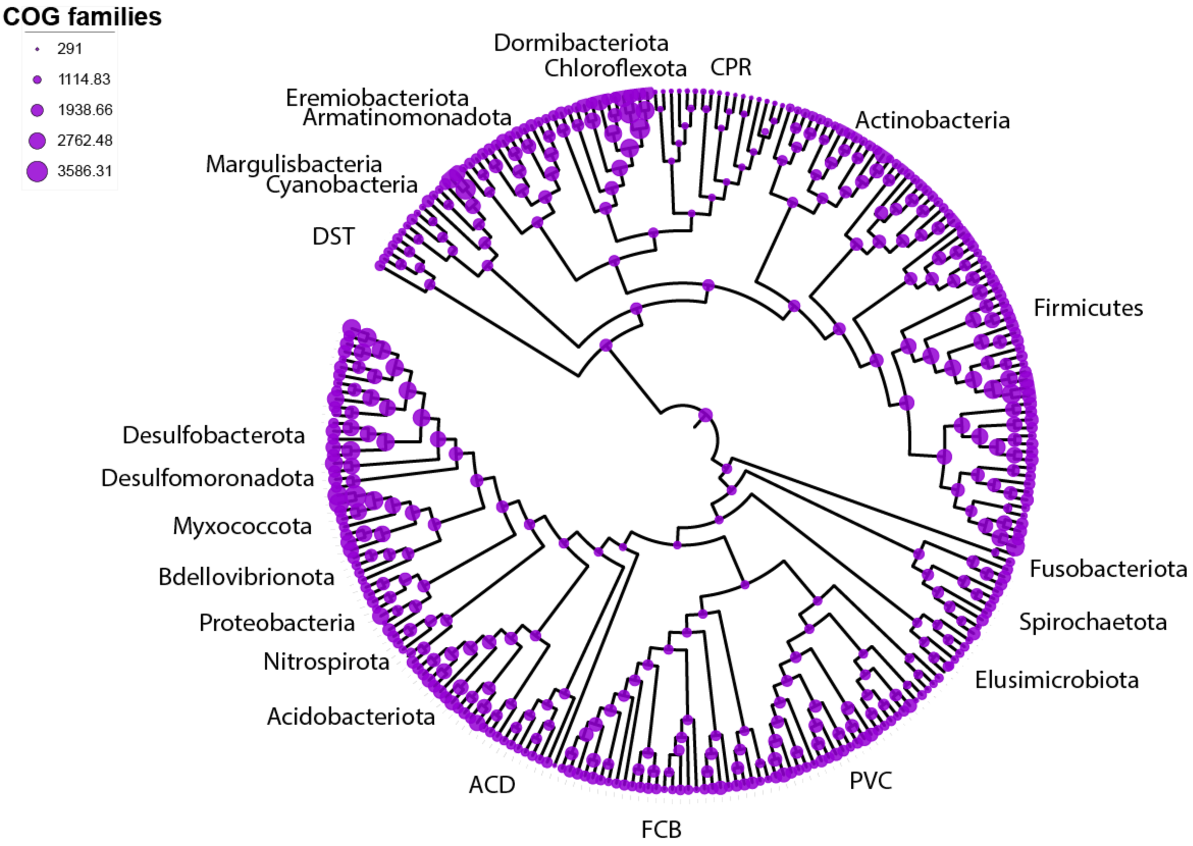
COG families
291
1114.83
1938.66
2762.48
3586.31
Dormibacteriota
Chloroflexota
CPR
Eremiobacteriota
Armatinomonadota
Actinobacteria
Margulisbacteria
Cyanobacteria
DST
Firmicutes
Desulfobacterota
Desulfomoronadota
Myxococcota
Bdellovibrionota
Proteobacteria
Nitrospirota
Acidobacteriota
Fusobacteriota
Spirochaetota
Elusimicrobiota
ACD
FCB
PVC

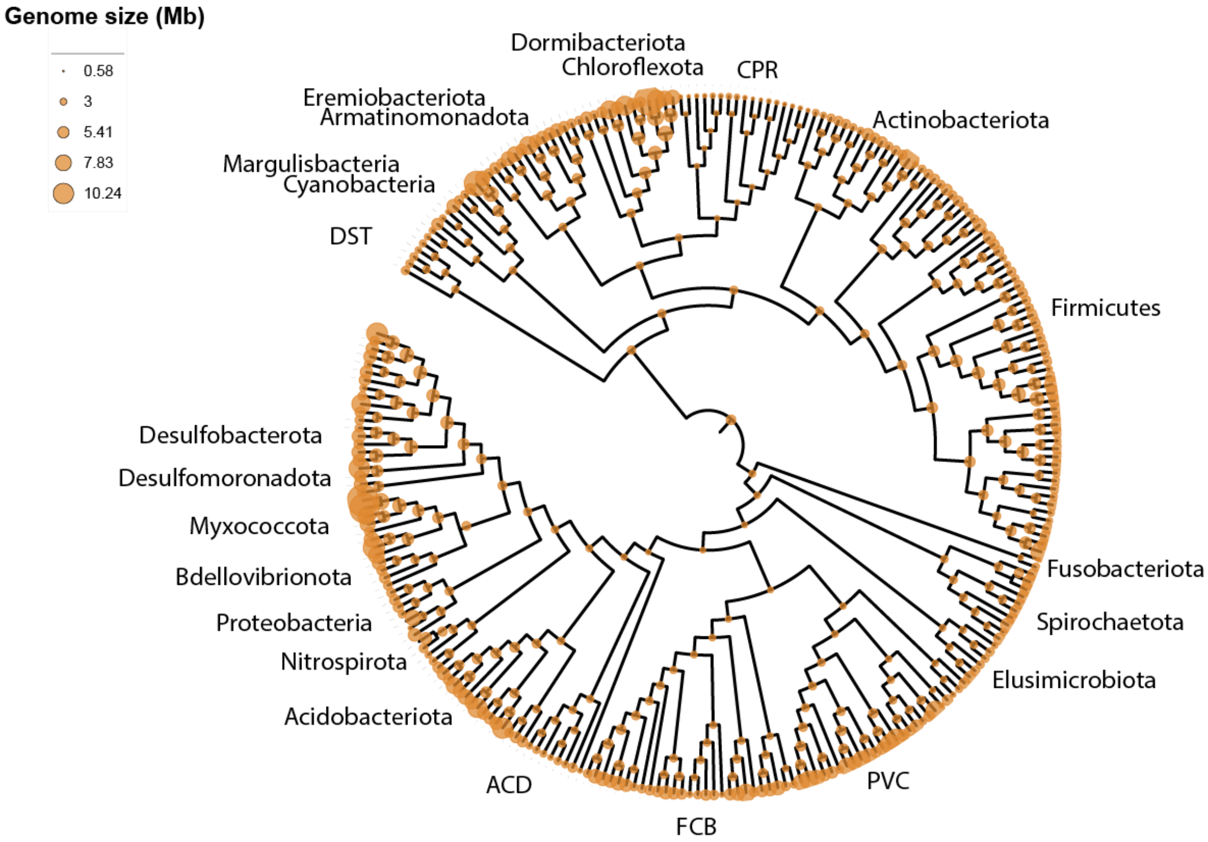
Genome size (Mb)
0.58
3
5.41
7.83
10.24
Dormibacteriota
Chloroflexota
CPR
Eremiobacteriota
Armatinomonadota
Actinobacteriota
Margulisbacteria
Cyanobacteria
DST
Firmicutes
Desulfobacterota
Desulfomoronadota
Myxococcota
Bdellovibrionota
Proteobacteria
Nitrospirota
Acidobacteriota
Fusobacteriota
Spirochaetota
Elusimicrobiota
ACD
FCB
PVC

**Extended Dated Figure 6: Evolution of COG family repertoires and inferred genome size over the bacterial tree.** (a) The inferred number of COG family members and (b) inferred genome size at each internal node of the tree. Genome sizes were predicted from the relationship between COG family members and genome size among extant Bacteria (LOESS regression). Circle diameter is proportional to family number or genome size. FCB are the Fibrobacterota, Chlorobiota, Bacteroides, and related lineages; PVC are the Planctomycetes, Verrucomicrobia, Chlamydiae, and related lineages; DST are the Deinococcous, Synergistota, and Thermatogota; ACD are Aquificota, Campylobacterota, and Deferribacterota; FA are Firmicutes and Actinobacteria. The figure depicts inferences for root 1 (as shown in Figure 1(b)); the data for all three roots are provided in GenomeSizeTable.tsv in the Extended Data Files.

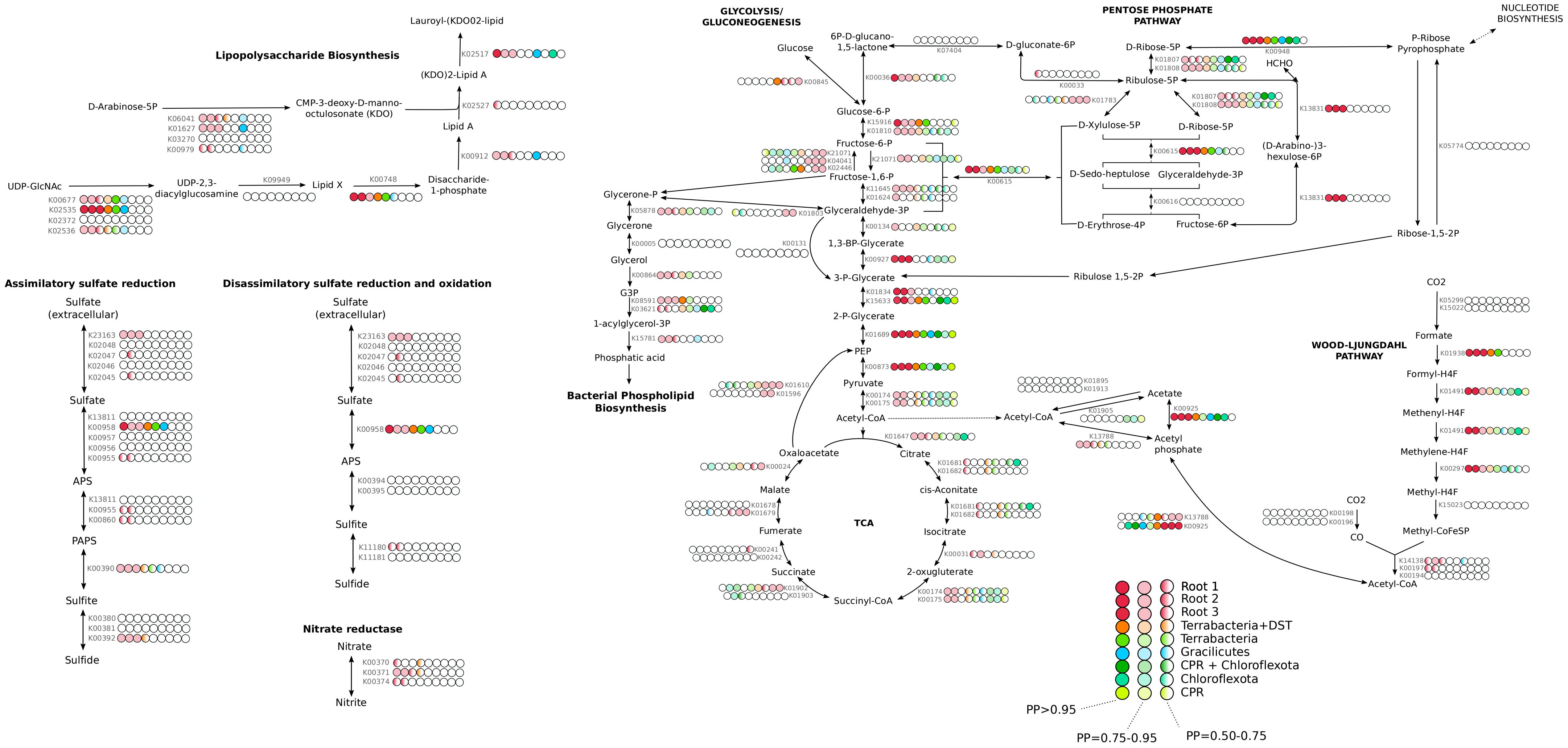
Lipopolysaccharide Biosynthesis
Lauroyl-(KDO02-lipid
(KDO)2-Lipid A
Lipid A
Disaccharide-1-phosphate
Lipid X
UDP-2,3-diacylglucosamine
UDP-GlcNAc
D-Arabinose-5P
CMP-3-deoxy-D-manno-octulosonate (KDO)
GLYCOLYSIS/ GLUCONEOGENESIS
PENTOSE PHOSPHATE PATHWAY
NUCLEOTIDE BIOSYNTHESIS
Glucose
6P-D-glucano-1,5-lactone
D-gluconate-6P
D-Ribose-5P
P-Ribose Pyrophosphate
HCHO
Ribulose-5P
Glucose-6-P
Fructose-6-P
Fructose-1,6-P
Glyceraldehyde-3P
D-Xylulose-5P
D-Sedo-heptulose
Glyceraldehyde-3P
D-Erythrose-4P
Fructose-6P
(D-Arabino-)3-hexulose-6P
Ribose-1,5-2P
Ribulose 1,5-2P
Glycerone-P
Glycerone
Glycerol
G3P
1-acylglycerol-3P
Phosphatic acid
Bacterial Phospholipid Biosynthesis
1,3-BP-Glycerate
3-P-Glycerate
2-P-Glycerate
PEP
Pyruvate
Acetyl-CoA
Acetyl-CoA
Acetate
Acetyl phosphate
Oxaloacetate
Citrate
Malate
cis-Aconitate
Fumerate
TCA
Isocitrate
Succinate
2-oxugluterate
Succinyl-CoA
WOOD-LJUNGDAHL PATHWAY
CO2
Formate
Formyl-H4F
Methenyl-H4F
Methylene-H4F
Methyl-H4F
Methyl-CoFeSP
CO2
CO
Acetyl-CoA
Root 1
Root 2
Root 3
Terrabacteria+DST
Terrabacteria
Gracilicutes
CPR + Chloroflexota
Chloroflexota
CPR
PP>0.95
PP=0.75-0.95
PP=0.50-0.75
Assimilatory sulfate reduction
Sulfate (extracellular)
Sulfate
APS
PAPS
Sulfite
Sulfide
Disassimilatory sulfate reduction and oxidation
Sulfate (extracellular)
Sulfate
APS
Sulfite
Sulfide
Nitrate reductase
Nitrate
Nitrite

**Extended Data Figure 7:** Metabolic map of the central metabolic pathways inferred in the last bacterial common ancestor (LBCA) and a selection of subsequent nodes. The reconstruction is based on genes that could be mapped to a given node with PP >0.5. The presence of a gene within a pathway is indicated as shown in the key. Annotations and PP values for KOs in this figure can be found in Supplementary Table 5. Annotations and PP values for all KOs can be found in Supplementary Table 4.

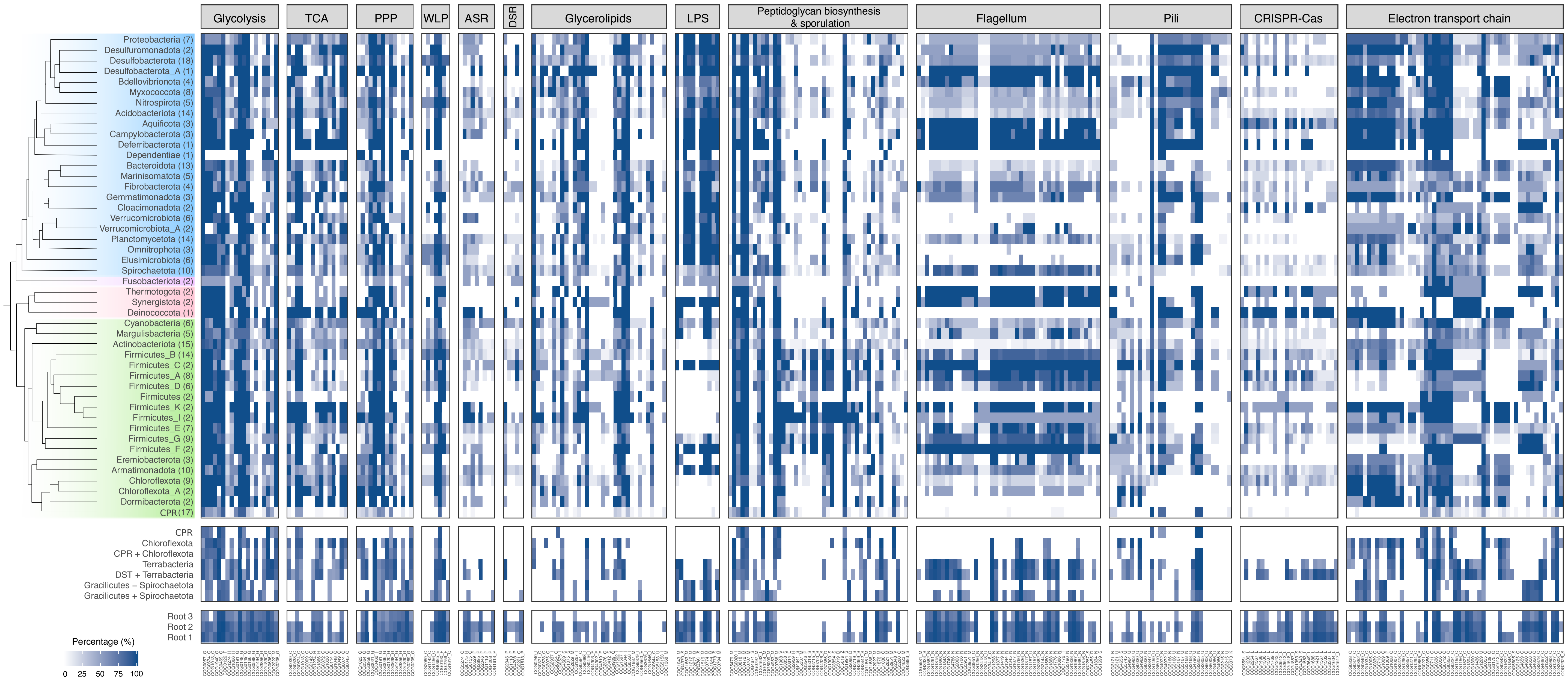

Glycolysis
TCA
PPP
WLP
ASR
DSR
Glycerolipids
LPS
Peptidoglycan biosynthesis & sporulation
Flagellum
Pili
CRISPR-Cas
Electron transport chain
Proteobacteria (7)
Desulfuromonadota (2)
Desulfobacterota (18)
Desulfobacterota_A (1)
Bdellovibrionota (4)
Myxococcota (8)
Nitrospirota (5)
Acidobacteriota (14)
Aquificota (3)
Campylobacterota (3)
Deferribacterota (1)
Dependentiae (1)
Bacteroidota (13)
Marinisomatota (5)
Fibrobacterota (4)
Gemmatimonadota (3)
Cloacimonadota (2)
Verrucomicrobiota (6)
Verrucomicrobiota_A (2)
Planctomycetota (14)
Omnitrophota (3)
Elusimicrobiota (6)
Spirochaetota (10)
Fusobacteriota (2)
Thermotogota (2)
Synergistota (2)
Deinococcota (1)
Cyanobacteria (6)
Margulisbacteria (5)
Actinobacteriota (15)
Firmicutes_B (14)
Firmicutes_C (2)
Firmicutes_A (8)
Firmicutes_D (6)
Firmicutes (2)
Firmicutes_K (2)
Firmicutes_I (2)
Firmicutes_E (7)
Firmicutes_G (9)
Firmicutes_F (2)
Eremiobacterota (3)
Armatimonadota (10)
Chloroflexota (9)
Chloroflexota_A (2)
Dormibacterota (2)
CPR (17)
CPR
Chloroflexota
CPR + Chloroflexota
Terrabacteria
DST + Terrabacteria
Gracilicutes − Spirochaetota
Gracilicutes + Spirochaetota
Root 3
Root 2
Root 1
Percentage (%)
0 25 50 75 100

**Extended Data Figure 8:** Distribution of COG families from key metabolic pathways inferred to the last bacterial common ancestor (LBCA). The occurrence of COG families in the taxa sampled in this study are represented as percentage presence across phylogenetic clusters (phylum) based on a presence/absence table. COG families inferred to the given nodes and the tree possible root positions (see Methods) are represented by corresponding PP values (PP>0.5). TCA=Tricarboxylic Acid Cycle, PPP=Pentose phosphate pathway, ASR=Assimilatory sulfate reduction, DSR=Dissimilatory sulfate reduction, LPS=Lipopolysaccharide. Supplementary Table 4 lists the PP values for all COG occurrences across roots, nodes, and tips and the metabolic genes featured in this plot can be found in Supplementary Table 5. A full heat map for all COGs, and additional heat maps by COGs category, can be found in Extended Data, in heatmaps.zip.

### Extended Data Tables

**Extended Data Table 1**: 63 orthologous genes used to infer the species tree, with those used in the outgroup rooting analysis indicated.

| KO number | Gene name | Annotation | Used in outgroup tree? |
|---|---|---|---|
| K03046 | rpoC | DNA-directed RNA polymerase subunit beta' | **y** |
| K03043 | rpoB | DNA-directed RNA polymerase subunit beta | |
| K02337 | dnaE | DNA polymerase III subunit alpha | **y** |
| K03070 | secA | Protein translocase subunit SecA | |
| K01873 | VARS, valS | Valine--tRNA ligase | |
| K02335 | polA | DNA polymerase I | **y** |
| K01872 | AARS, alaS | Alanine tRNA ligase | |
| K02469 | gyrA | DNA gyrase subunit A | |
| K00962 | pnp, PNPT1 | Polyribonucleotide nucleotidyltransferase | **y** |
| K02355 | fusA, GFM, EFG | Translation elongation factor G | **y** |
| K01972 | E6.5.1.2, ligA, ligB | DNA ligase NAD | |
| K03702 | uvrB | Excinuclease ABC subunit B | |
| K02470 | gyrB | DNA gyrase subunit B | |
| K04077 | groEL, HSPD1 | Molecular chaperone GroEL | |
| K01937 | pyrG, CTPS | CTP synthase | **y** |
| K02313 | dnaA | Chromosomal replication initiator protein | |
| K02314 | dnaB | Replicative DNA helicase | |
| K02433 | gatA, QRSL1 | aspartyl-tRNA(Asn)/glutamyl-tRNA(Gln) amidotransferase subunit A | |
| K03076 | secY | Protein translocase subunit SecY | **y** |

| | | | |
|---|---|---|---|
| K04485 | radA, sms | DNA repair protein RadA/Sms | |
| K02112 | ATPF1B, atpD | F-type H+/Na+-transporting ATPase subunit beta | **y** |
| K03590 | ftsA | Cell division protein FtsA | |
| K02358 | tuf, TUFM | Elongation factor Tu | **y** |
| K06942 | ychF | Redox Regulated ATPase YchF | |
| K00927 | PGK, pgk | Phosphoglycerate kinase | |
| K01889 | FARSA, pheS | Phenylalanine--tRNA ligase alpha subunit | |
| K03551 | ruvB | Holliday junction branch migration DNA helicase RuvB | **y** |
| K04485 | radA, sms | DNA recombination repair protein RecA | **y** |
| K02835 | prfA, MTRF1, MRF1 | Peptide chain release factor 1 | |
| K02886 | RP-L2, MRPL2, rplB | 50S ribosomal protein L2 | **y** |
| K01803 | TPI, tpiA | Triose phosphate isomerase | **y** |
| K03438 | mraW, rsmH | 16S rRNA (cytosine1402-N4)-methyltransferase | |
| K00554 | trmD | tRNA (guanine37-N1)-methyltransferase | |
| K02863 | RP-L1, MRPL1, rplA | 50S ribosomal protein L1 | **y** |
| K03685 | rnc, DROSHA, RNT1 | Ribonuclease III | |
| K02967 | RP-S2, MRPS2, rpsB | 30S ribosomal protein S2 | **y** |
| K02982 | RP-S3, rpsC | 30S ribosomal protein S3 | |
| K02906 | RP-L3, MRPL3, rplC | 50S ribosomal protein L3 | **y** |
| K03470 | rnhB | Ribonuclease HII | |
| K01358 | clpP, CLPP | ATP dependent Clp protease proteolytic subunit | |
| K06187 | recR | Recombination protein RecR | |
| K15034 | yaeJ | Aminoacyl tRNA hydrolase, ribosome-associated protein | |
| K02931 | RP-L5, MRPL5, rplE | 50S ribosomal protein L5 | **y** |
| K02933 | RP-L6, MRPL6, rplF | 50S ribosomal protein L6 | **y** |
| K02601 | nusG | Transcription termination antitermination protein NusG | |
| K02988 | RP-S5, MRPS5, rpsE | 30S ribosomal protein S5 | **y** |
| K02992 | RP-S7, MRPS7, rpsG | 30S ribosomal protein S7 | **y** |
| K03664 | smpB | SsrA binding protein | |
| K02838 | frr, MRRF, RRF | Ribosome recycling factor | |
| K02867 | RP-L11, MRPL11, rplK | 50S ribosomal protein L11 | |
| K02878 | RP-L16, MRPL16, rplP | 50S ribosomal protein L16 | **y** |
| K02871 | RP-L13, MRPL13, rplM | 50S ribosomal protein L13 | **y** |
| K02994 | RP-S8, rpsH | 30S ribosomal protein S8 | **y** |

| | | | |
|---|---|---|---|
| K02948 | RP-S11, MRPS11, rpsK | 30S ribosomal protein S11 | **y** |
| K02952 | RP-S13, rpsM | 30S ribosomal protein S13 | **y** |
| K02935 | RP-L7, MRPL12, rplL | 50S ribosomal protein L7/12 | |
| K02996 | RP-S9, MRPS9, rpsI | 30S ribosomal protein S9 | |
| K02874 | RP-L14, MRPL14, rplN | 50S ribosomal protein L14 | **y** |
| K02887 | RP-L20, MRPL20, rplT | 50S ribosomal protein L20 | |
| K02946 | RP-S10, MRPS10, rpsJ | 30S ribosomal protein S10 | **y** |
| K02965 | RP-S19, rpsS | 30S ribosomal protein S19 | **y** |
| K02956 | RP-S15, MRPS15, rpsO | 30S ribosomal protein S15 | |
| K02518 | infA | Translation initiation factor IF 1 | |

**Extended Data Table 2**: Support for published hypotheses using outgroup rooting.

| **Root hypothesis** | **log-likelihood difference to ML** | ***p*-value** | **Study** |
|---|---|---|---|
| Observed outgroup root (Supplementary Fig. 5) | 0 | 0.71 | This study (ML tree) |
| Between Gracilicutes and Terrabacteria | -7.6 | 0.55 | This study (ALE root, see below) |
| Thermotoga/Synergistes/Deinococcus basal* | -11.5 | 0.48 | ? |
| Chloroflexi basal | -11.6 | 0.46 | ? |
| Planctomycetes basal | -13.4 | 0.47 | ? |
| DPANN basal within archaeal outgroup | -19.9 | 0.41 | [1,3] |
| CPR basal | -20.4 | 0.35 | [4,5] |
| Between Firmicutes and Actinobacteria | -26.8 | 0.36 | ? |
| Fusobacteria basal | -27.3 | 0.32 | ? |

**Extended Data Table 3**: **Support for published rooting hypotheses from our outgroup-free analyses.** *Our unrooted topology was incompatible with some published hypotheses, including a clade of Thermotogales and Aquificales at the root[6,7].

| **Root** | ***p*-value** | **Study** |
|---|---|---|
| CPR basal | 2e-04 | Hug et al. (2016), Zhu et al. (2019) |
| Chloroflexi basal | 1e-41 | Cavalier-Smith (2006) |

| | | |
|---|---|---|
| Between Firmicutes and Actinobacteria | 9e-05 | Lake et al. (2009) |
| Thermotoga/Synergistes/Deinococcus basal* | 0.004 | |
| Planctomycetes basal | 2e-26 | Brochier and Philippe (2002) |

### Supplementary Tables

**Supplementary Table 1: AU *p*-values for all tested roots in ALE analysis (Excel-formatted spreadsheet)**

**Supplementary Table 2: AU-test results for an ALE root analysis using 3595 COG families.**

| Root name | LLs | AU |
|---|---|---|
| Fusobacteria root (398) | -7.3 | 0.589 |
| Fusobacteria on Terrabacteria side (527) | 7.3 | 0.519 |
| Fusobacteria on Gracillicutes side (528) | 13.6 | 0.432 |
| Fusobacteria and Spirochaeta on Terrabacteria side (520) | 32.1 | 0.251 |
| DST root (464) | 103.7 | 0.008 |
| Cyanobacteria on Gracillicutes side (517) | 215.8 | 1e-09 |
| Dormi/Chloroflexi+CPR (510) | 372.2 | 7e-06 |
| CPR root (496) | 425.2 | 2e-67 |
| Dormi/Chloroflexi (505) | 630.7 | 5e-05 |
| Omnitrophota/Verrucomicrobia/Planctomycetes (492) | 674.4 | 2e-04 |
| Fibrobacteria/Bacteroidetes/Marinisomatota (511) | 1000.7 | 3e-71 |

**Supplementary Table 3: Singleton support (the number of genes that evolve vertically from one end of a branch to the other) on the credible set of rooted trees.** Root numbers correspond to the three root branches depicted in Figure 1(b).

| **Root branch** | **1** | **2** | **3** |
|---|---|---|---|
| Median singleton support per branch | 999.09 | 998.47 | 999.21 |

| | | | |
|---|---|---|---|
| Singleton support for branches subtending the root | 98.259, 140.45 | 91.56, 151.09 | 95.25,117.16 |
| Mean verticality | 0.68136 | 0.68137 | 0.6818 |

**Supplementary Table 4: Protein family annotations (COG and KO) and root presence posterior probabilities (PPs) for all 3723 gene families under all three branches in the root region (Excel-formatted spreadsheet).**

**Supplementary Table 5: Protein family annotations (COG and KO) and root presence posterior probabilities (PPs) for key pathways used in reconstruction (Excel-formatted spreadsheet).**

**Supplementary Table 6: COG families lost on the CPR stem (Excel-formatted spreadsheet).**

**Supplementary Table 7: Mean verticality V/(V+T) by COG functional category.**

| COG | Annotation | Number of families | Median verticality | Mean verticality |
|---|---|---|---|---|
| **V** | Defense mechanisms | 32 | 0.5280122078075817 | 0.5638668252634967 |
| **T** | Signal transduction | 88 | 0.5799451471715107 | 0.604432180596873 |
| **G** | Carbohydrate | 186 | 0.5913637562346645 | 0.6015642843633491 |
| **Q** | Secondary metabolites | 73 | 0.5964080412431355 | 0.6073596199333191 |
| **L** | Replication | 179 | 0.5987938232160366 | 0.6150912826158655 |
| **P** | Inorganic ion | 199 | 0.5988930441950502 | 0.6115949584832507 |
| **O** | Post-translational modification | 123 | 0.6013500360389593 | 0.6131412846715851 |
| **K** | Transcription | 131 | 0.6078333513305463 | 0.6340799581816328 |
| **I** | Lipid | 77 | 0.6124862586859439 | 0.6216075914824997 |
| **C** | Energy | 239 | 0.6145542409023125 | 0.6275371657010402 |
| **M** | Cell wall/membrane | 141 | 0.6155364029228826 | 0.625358402235949 |
| **H** | Coenzyme | 165 | 0.6233926976132386 | 0.6295188636764987 |
| **E** | Amino acid | 226 | 0.6235372737809106 | 0.631566055802421 |
| **F** | Nucleotide | 102 | 0.64234381501306 | 0.6370022274662093 |
| **N** | Cell motility | 70 | 0.6825519330347292 | 0.6801963045783017 |
| **D** | Cell cycle | 45 | 0.6849512300407962 | 0.6905029550824039 |

| | | | | |
|---|---|---|---|---|
| **U** | Intracellular trafficking | 83 | 0.6854327621149885 | 0.6880748036306573 |
| **J** | Translation | 177 | 0.6906677776226816 | 0.6870530339985472 |

**Supplementary Table 8: Number of taxa sampled from each clade in the GTDB-independent analysis.**

| Clade | No. of taxa sampled |
|---|---|
| Firmicutes | 25 |
| Actinobacteriota+Cyanobacteria+Chloroflexota | 35 |
| CPR | 125 |
| FCB+PVC+Elusimicrobiota | 35 |
| Proteobacteria | 50 |
| Deltaproteobacteria+Nitrospirota+Acidobacterota+Aquificota | 30 |
| FASST+environmental lineages | 25 |
| New Phyla (Parks et 2018) | 17 |

**Supplementary Table 9: Estimated root origination rates and root presences by COG functional category.**

| COG category | Number at root (flat prior) | Number at root (ML prior, PP >= 0.95) | Number at root (ML prior, PP >= 0.8) | Number of families in category | ML O_R |
|---|---|---|---|---|---|
| J | 3 | 125 | 157 | 177 | 6739.434 |
| F | 1 | 57 | 89 | 102 | 5294.183 |
| L | 4 | 42 | 100 | 179 | 1468.033 |
| H | 3 | 26 | 85 | 165 | 1393.54 |
| - | 0 | 0 | 1 | 2 | 1290.933 |
| N | 1 | 18 | 38 | 70 | 1270.758 |
| E | 2 | 45 | 116 | 226 | 1146.195 |
| D | 2 | 7 | 16 | 45 | 850.0477 |
| P | 4 | 16 | 64 | 199 | 650.1995 |
| M | 1 | 15 | 49 | 141 | 625.6504 |
| C | 4 | 22 | 57 | 239 | 581.1914 |

| | | | | | |
|---|---|---|---|---|---|
| G | 3 | 11 | 43 | 186 | 513.1411 |
| I | 0 | 1 | 12 | 77 | 457.9165 |
| U | 1 | 3 | 12 | 83 | 364.7495 |
| K | 1 | 3 | 10 | 131 | 260.9843 |
| O | 0 | 1 | 9 | 123 | 256.3592 |
| S | 6 | 9 | 69 | 1374 | 163.0099 |
| B | 0 | 0 | 1 | 5 | 126.1288 |
| T | 0 | 1 | 3 | 88 | 125.0658 |
| V | 0 | 0 | 0 | 32 | 76.72498 |
| Q | 0 | 0 | 0 | 73 | 1.196214 |
| A | 0 | 0 | 0 | 4 | 1 |
| Z | 0 | 0 | 0 | 2 | 1 |

### Extended Data Files

The following files have been deposited at FigShare, DOI: 10.6084/m9.figshare.12651074.

**SpeciesTable.tsv** - Table containing the mapping of the short names (nX) to GTDB accessions and taxonomy.

**RootTable.tsv -** Table containing the AU values, the likelihood difference, and the genomes on both sides of the root (separated by ;;;).

**VerticalityTable.tsv** - Table containing the verticality of every COG family.

**ReconciliationsMCL.zip** - Folder containing the reconciliations files for the 3 possible roots, using the MCL families.

**ReconciliationsCOGs.zip** - Folder containing the reconciliations files for the 3 possible roots, using the COG families.

**RelativeDating.zip -** Folder containing the highly supported constraints for the 3 possible rooted species trees and the 1000 sampled orders compatible with those constraints.

**SpeciesTrees.zip -** Folder containing the three rooted species trees in Newick format.

**GenomeSizeTable.tsv -** Table containing the results from the LOESS regression analysis of COG family members against genome size.

**Orthologues_for_species_tree.zip** - Alignments for the orthologues used to compute the species tree for the focal analysis.

**FocalTreeAnalysis.zip** - Concatenate and Newick tree for the focal tree analysis.

**CompositionallyStrippedAnalysis.zip -** Alignments used to infer the compositionally-stripped trees and the trees in Newick format.

**OutgroupAnalysis.zip** - Alignment and tree in Newick format used in the outgroup analysis.

**Code.zip -** Folder containing the following scripts:

**O_R_Optimization.py** - Python code to estimate the posterior probability of presence at the root for each gene family.

**mc_explorer.py** - Python script that computes randomly generated time orders compatible with a set of constraints.

**large_COGs.txt -** List of COGs for which conditional clade probabilities could not be computed due to large size.

**heatmaps.zip -** Supplementary heat maps, including full heat map of all COGs and heat maps for each COG category.

**HeatmapScripts.zip -** Folder contains the following scripts:

**PPs_table_parsing.R** - R script for generating heat maps.

**BRooting_Annotation_workflow.sh -** Contains workflow for protein and protein family functional annotations, KO assignment to COG families, and parsing of raw PP data into a readable PP table (Supplementary Table 4)

**HeatMapMappingFiles.zip -** Folder containing mapping files for producing heatmaps:

**COGs_of_interest_metabolism_vs3.txt** - contains all COGs associated with Metabolism processes.

**COGs_of_interest_MetMap_vs3-2.txt** - contains the set of COGs used for metabolic reconstruction.

**ShortName_to_taxa_mapping_file_vs5.txt** - short name-to-taxa mapping file built from SpeciesTable.tsv.

**V4_all_pp_roots_and_tips_cln.txt** - contains the final PP table used for the construction of heat maps derived from Supplementary Table 4.

### Supplementary Methods and Discussion

#### Supplementary Methods

##### Taxon sampling for focal analysis

To obtain a representative taxon sampling from across known bacterial diversity, we sampled taxa according to the classification provided by the Genome Taxonomy Database (GTDB r89)[8]. We sampled 265 genomes from the GTDB as follows. First, we filtered out the genomes with Quality < 0.75 (Quality is defined as Completeness - (5*Contamination)[9]), and filtered out all phyla subsequently left with fewer than 10 species. Genomes were sampled from the remaining taxa on a per-class basis: for classes containing a single order, the genome with the highest quality score was sampled; for classes containing multiple orders, the highest quality genome from each of two randomly chosen orders was sampled. This protocol ensured that every class in the GTDB is represented in the final tree. We then manually added the genome of *Gloeomargarita litophora* given its importance in constraining the phylogeny and timing of chloroplast evolution. The list of genomes can be found in the Data Supplement, at SpeciesTable.tsv.

##### Gene family clustering and ALE analysis

We used the protein annotations provided by GTDB, which were originally obtained using Prodigal. To infer homologous gene families for ALE inference, we performed an all vs all similarity search using Diamond[10] with an E-value threshold of $<10^{-7}$ to avoid distant hits and k = 0 to report all the relevant hits. Current clustering methods are not consummate and the parameters that determine the granularity of clustering do not have a direct biological motivation. Setting the value of the MCL inflation parameter therefore involves a trade-off between inferring large, inclusive clusters that will contain false positives (sequences that are not part of the real gene family) and small, conservative clusters that may divide real gene families into several subclusters. An additional practical concern for phylogenomics is that overly large clusters can align poorly and result in low-quality single protein trees. In our rooting analysis, we experimented with a range of values for the mcl inflation parameter, and chose 1.2 because the clusters were inclusive without a substantial reduction in post-masking alignment length compared to more granular settings.

Clustering using MCL[11] with an inflation parameter of 1.2 resulted in 186,827 gene families and a total of 11,765 families with 4 or more sequences. We aligned the 11,765 gene families using MAFFT[12] (with the --auto option) and filtered with BMGE[13] (using bmge -t AA -m BLOSUM30) After filtering, 260 alignments contained no high-quality columns and were discarded. We filtered out sequences containing more than 80% of gaps to produce the final set of alignments. We also discarded all alignments with less than 30 columns, leaving a total of 11,272 families. The gene trees were computed using IQ-TREE v 1.6.10 using the following command:

```
iqtree -m TEST -s FAMXXX.faa.aln.trimmed -bb 10000 -wbtl -nt AUTO -madd
LG4X,LG4M,LG+C10,LG+C20,LG+C30,LG+C40,LG+C50,LG+C60,C10,C20,C30,C40,C50,C60.
```

Conditional clade probabilities (CCPs) were computed using ALEobserve and the resulting ALE files were reconciled with the species tree. Loss rates were corrected by genome completeness, estimated using CheckM[14]. We tested 62 roots (Extended Data, RootTable.tsv).

**GTDB-independent analysis**

To sample representative bacterial taxa independently of the GTDB, we began with the bacterial portion of a recent global analysis of the tree of life[4]. We inferred a tree of the bacterial portion of the concatenate under the LG+G4+F model in IQ-Tree. We divided the tree into 7 major bacterial clades based on a literature search (Supplementary Table 8) and additional environmental lineages with branch length diversity comparable to the known groups. For each group defined in this way, we manually subsampled taxa so as to maintain genetic diversity, while avoiding overly long or short branches. We selected 342 species, comprising 200 'classic' bacteria, 125 CPR bacteria and one bacterial genome respectively from each of the 17 new phyla described by[9]. We used the same marker gene set as in the focal analysis. A species tree was inferred in IQ-Tree using the LG+C20+G4 model with PMSF[15]. Additional trees were inferred in PhyloBayes under the CAT+GTR+G4 model using a recoded alignment using the four-category scheme of Susko and Roger[16], and under the multispecies coalescent model in ASTRAL[17]. To infer homologous gene families, we used the same pipeline as that used in the focal analysis. This resulted in 15,592 gene families with 4 or more sequences, which were analysed as in the focal analysis. The unrooted species trees are congruent with each other, except in the placement of a small group of phyla comprising Fusobacteriota, Aquificota, Synergistota, Spirochaetota and Thermtogota ("FASST"). These taxa are resolved in different positions in each of the three unrooted topologies, and are found to be monophyletic in the tree inferred from the recoded alignment. Similarly to the focal analysis, the ALE analyses yielded two root positions that could not be rejected (AU test, $p > 0.05$), summarised in Extended Data Figure 3. Both of these rooted phylogenies are congruent with that of the focal (GTDB) analysis, with Terrabacteria and Gracilicutes on either side of the root, with the only differences being in the placement of the FASST taxa. In the focal analysis, as well as the LG+PMSF+G4 and ASTRAL trees inferred as part of the GTDB-independent analysis, FASST were not recovered as a monophyletic group.

**Ancestral gene content and metabolic reconstruction**

*COG gene families for ancestral gene content reconstruction*

We built a set of gene families based on the COG[18] database for ancestral functional inference. To do so, we annotated each genome in the dataset using eggNOG-mapper v2[19], then clustered proteins into families based on their COG annotations. For proteins annotated with more than one COG category (8% of proteins), we included the protein in both COG families. This resulted in 4256 COG families, of which 3723 had 4 or more sequences. COG families are ideal for ancestral reconstruction because they comprise all of the sequences on extant genomes that can be annotated with a given unambiguous function from the COG ontology. In addition, the hierarchical nature of the COG classification (comprising gene family annotations nested within 23 broader functional categories) enabled us to explicitly model the different evolutionary ages of gene functional classes as part of the analysis, by using category-specific root origination priors (see below).

Our COG families are useful for functional reconstruction, but are perhaps less well suited for investigating other aspects of bacterial evolution because they are constructed only from proteins that could be annotated with eggNOG-mapper. By contrast, MCL families represent --- within the limitations of the clustering approach, as discussed above --- an unbiased view of gene family diversity for the set of genomes we analyzed. We therefore base analyses other than those regarding the functional annotation of LBCA on the MCL families. However, since gene clustering methods are not consummate and each has strengths and weaknesses, we also investigated the root signal from the COG families. This analysis resulted in a root region of four adjacent branches, comprising the root region from the focal analysis (3 branches) plus one additional branch, in which Spirochaetota branched on the Terrabacteria side of the root (Supplementary Table 2). This slightly expanded root region is likely due to the reduced resolution of the smaller set of COG families in comparison to the full analysis.

*Root gene mapping approach*

To estimate root presence posterior probabilities (PPs) for each gene family for each of the three supported roots, we first estimated the root origination prior (O_R) by maximum likelihood, finding the O_R value that maximises the total reconciliation likelihood summed over all gene families. We then used the global ML O_R value to calculate the root presence posterior probabilities for each family; that is, the probability that one or more copies of a given gene family were present at the root, given the ML O_R value. These indicated that families with different functions varied widely in terms of root presence probability, in agreement with established theory[20]; for example, proteins involved in translation (J) had the highest root presence probabilities among the functional classes investigated (Supplementary Table 9). We therefore estimated root origination rates independently for each of the 23 COG functional categories, and used these rates to estimate the posterior probability of presence at the root node for each gene family. Note that, for all nodes of the tree (including the root nodes), we additionally estimated PPs directly from the sampled reconciliations. Python code implementing this procedure is provided in the Extended Data Files at Code/O_R_Optimization.py.

Initial gene content and metabolic inferences at a particular node were based on gene families with a posterior presence probability (PP) of >0.95 at that node. This approach is conservative and can result in a range of PP values for different proteins within a metabolic pathway. Therefore, we manually investigated the PPs of pathways discussed in this manuscript and inferred the presence of specific pathways or functional modules if the majority of its components were found with PP >0.5, as described in the main text and Supplemental Discussion, Figure 4 and Extended Data Figure 7.

*Impact of root branch on LBCA gene content*

The credible set of root branches from the ALE analysis comprised three adjacent branches at the centre of the tree (Figure 1b). The difference between these three root positions relates to the placement of Fusobacteria, either as the root branch or as the most basal split on either the Gracilicutes or Terrabacteria+DST “sides” of the rooted tree. We therefore estimated root PPs for COG families on all three branches; Supplementary Table 4 provides root PPs under all three roots and indicates when genes were present in 1, 2, or all 3 candidate root positions.

### Supplementary Discussion

*Performance of ALE for species tree rooting*

The ability of the ALEml_undated algorithm to infer the correct gene tree root in the presence of gene duplications, transfers and losses was previously investigated using simulations[1]. Briefly, gene families were simulated on a rooted species tree using a continuous-time ODTL process (that is, a more complex model of genome evolution than that implemented in ALEml_undated), and ALEml_undated was used to estimate the root from subsamples of the simulated families. The maximum likelihood root according to ALE was the correct root in 95/100 replicates, and the log likelihood of alternative roots decreased with nodal distance from the correct root (as observed in our empirical data, see Figure 1). In the remaining 5 cases, the maximum likelihood root was one branch away from the true root. Analysis of empirical data suggested that ALE root inferences are robust to (that is, consistent across) subsets of the data that vary in terms of the rate of horizontal gene transfer or species representation in gene families[1]. These properties make the ALE approach appropriate for inferring the root of Bacteria.

*Performance of ALE for inferring rooted gene trees*

More broadly, species tree-aware phylogenetic methods (such as ALE) have been shown to be of use in fixing gene tree errors[21], and for ancestral state[1] and protein[22] inference. The additional power of these methods derives from the use of information in the species tree to decide between gene trees that are statistically equivalent from the point of view of the phylogenetic likelihood. Recently, a study of gene tree rooting performance suggested that a parsimony-based, species tree unaware DTL method (RANGER-DTL) provided more accurate gene tree root estimates than species tree-aware, probabilistic methods such as ALE and GeneRax. Gene tree rooting accuracy is not directly related to species tree rooting accuracy, because the information on the species tree root in ALE derives from finding the rooted species tree that maximises the sum of reconciliation likelihoods across gene families. We nevertheless decided to investigate, in order to understand which properties of the available methods contribute to, and detract from, rooting accuracy more generally.

To investigate, we re-analysed the data from Figure 5 of Morel et al.[23], who simulated sequence alignments on known rooted gene trees. This setup differs from that of the original study[24] in that, where possible, we start directly from the alignment and not from independently reconstructed gene trees. We chose this simulation setup because empirical analyses typically proceed from sequence alignments to inferred trees and then roots. We inferred rooted gene trees roots using ALE, GeneRax[23] (a species-tree *aware* method that maximises a joint reconciliation and phylogenetic likelihood to infer rooted gene trees), TreeRecs (a species-tree aware method that implements a parsimony approach to DTL to infer rooted gene trees), RANGER-DTL[25] (a species-tree *unaware* method that implements a DTL parsimony approach to root input gene trees), MAD[26] (a species-tree unaware method that roots input gene trees using minimal ancestor deviation). To quantify gene tree rooting accuracy, we plotted the rooted RF score between the inferred and true rooted gene trees; this provides an interpretable measure of rooting accuracy even when the true root bipartition does not appear in the inferred gene tree, as demonstrated recently[24]. The results (Supplementary Figure 1) indicate that probabilistic species tree-aware methods (GeneRax and ALE) provide the most accurate gene tree root inferences among the methods compared, even when the model used to infer the gene tree is misspecified (that is, when LG is used for simulation and WAG for inference; in particular, even when gene trees are inferred with a misspecified substitution model, ALE gene tree rooting (mean rooted-RF = 0.258) is significantly more accurate than RANGER-DTL (mean rooted-RF = 0.322, $p < 10^{-15}$ Welch Two Sample t-test) or MAD (mean rooted-RF = 0.306, $p < 10^{-10}$ Welch Two Sample t-test) when the latter methods are provided with input gene trees obtained using the true substitution model.

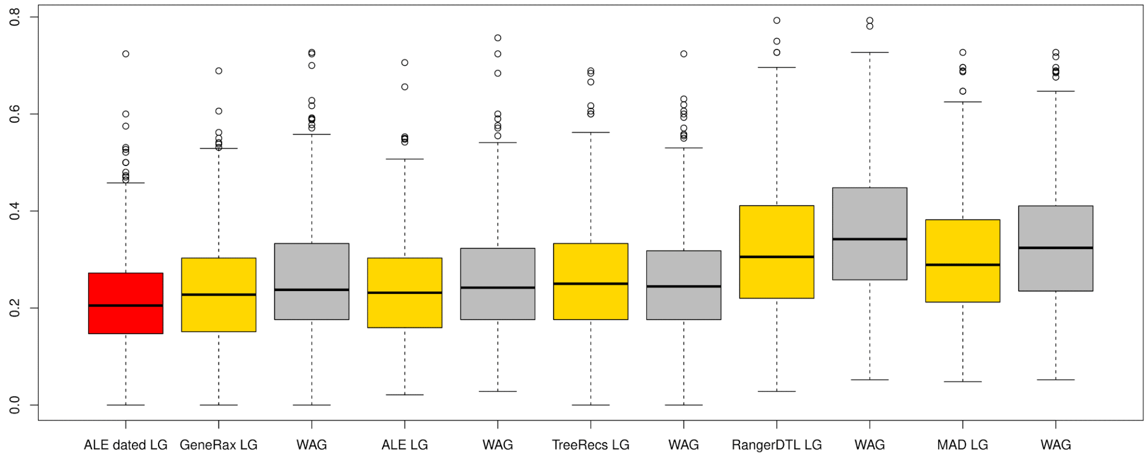


**Supplementary Figure 1:** Accuracy of gene tree rooting methods. Species tree aware methods (GeneRax, ALE, TreeRecs) are the most accurate, even when gene trees are inferred with a misspecified substitution model. Among species tree unaware methods, MAD outperforms the parsimony-based DTL method RANGER-DTL.

#### Ancestral reconstruction of the proteome of LBCA

*Information processing, cell division and signaling*

For the following discussion regarding metabolic reconstructions, we refer to the three branches in the root region as Root 1, Root 2, and Root 3 respectively, as shown in Figure 1b. In what follows, we provide probability ranges across the three branches of the root region for the presence of key genes. One caveat of our analyses is that the method has limited power to distinguish between ancestral presence in LBCA as opposed to origin on an early descendant branch followed by gene transfer; this may contribute to the mapping of some combinations of gene families to LBCA that are likely to be ancient but, on physiological grounds, are unlikely to have coexisted in a single cell (see below).

A large number of genes that can be mapped back to LBCA are involved in informational processing and storage machineries such as translation, transcription and replication in all three roots with PPs above 0.5, with many being highly supported (PP>0.95). This includes the majority of ribosomal proteins, tRNA synthetases and genes involved in their biosynthesis. We also recovered DNA polymerase I, III and IV, DNA topoisomerase type I, and DNA ligase. Additionally many exonucleases, endonucleases and ribonucleases are recovered, as well genes for base excision repair, nucleotide excision repair, mismatch repair and homologous recombination (Supplementary Table 4).

Furthermore, we detected many genes for cellular processes and signalling, including cell division, signal transduction, membrane transport, intracellular trafficking, chemotaxis and cellular mobility. For example, the PP for the presence of the cell division proteins FtsZ, FtsQ and FtsA (K03531, K03589, and K03590) were >0.9 in each root. A small number of proteins involved in signal transduction, i.e. two-component regulatory systems, had a PP>0.95, with many more genes recovered with a PP>0.5. Additionally, we recover genes for components of 16 ABC transporters across all three roots, with 23 recovered in both Roots 1 and 2 (PP >0.5). We also recover evidence for the bacterial secretion system, with four proteins of Sec system having a PP>0.8 across all three roots. Two of these, SecY (K03076) and SecD/F (K12257) had a PP>0.95 in roots 1 and 2. Additionally, the GspD (K02453) and GspJ (K02459) subunits of secretion system II were recovered with a PP>0.8 in all three roots, or with a PP >0.5 in Root 1 and 2, respectively (Supplementary Table 4).

*Cell envelope and motility*

One interesting aspect of prokaryotic evolution is the so-called ‘lipid divide’[27] Typically, Archaea have G1P phospholipids with ether bonds and isoprenoid chains, which are often membrane spanning [28]. Bacteria, on the other hand, possess G3P phospholipids, typically with ester bonds and fatty-acid chains, that form bilayers[28]. While several recent studies indicate the existence of lipids with mixed characteristics, their provenance and the mechanisms by which they are synthesised are currently unclear[29–32]. Nonetheless, such discoveries have led to the questioning of the lipid divide. Genes encoding components of archaeal lipids have been found to be widespread in Bacteria[33,34] and genes encoding components for bacterial lipids are found in Archaea, although to a lesser extent[34]. Furthermore, a recent study demonstrated that *E. coli* engineered to have a stable hybrid heterochiral lipid membrane do not experience any change in growth rate[35]. While, phylogenetic analyses suggest that the archaeal pathway may predate the bacterial one[34], there was no significant support for the presence of archaeal lipid biosynthesis genes in LBCA. In particular, genes coding for the enzymes that determine the phospholipid stereochemistry did not have PPs above a threshold of 0.5 (see Supplementary Table 5), though a gene coding for glycerol-3-phosphate (K00111) might have been present in Root 1 (PP=0.49). We do, however, recover glycerol kinase (GlpK), which can synthesise G3P form glycerol (K00864, PP=0.9/0.87/0.73). Furthermore, our analyses suggest the presence of PlsY (K08591, PP=0.94/0.92/0.82) and PlsX (K03621, PP=0.71/0.68/0.38), which attach the first fatty acid chain to G3P in many Bacteria (some species alternatively use PlsB, which we do not recover

PP>0.5). We also recover a putative PlsC (K15781, PP=0.86/0.83/0.64), which attaches the second fatty-acid side-chain. Our inferences therefore suggest that LBCA had bacterial phospholipid membranes, while being unable to synthesize archaeal lipids.

Bacteria have classically been divided into Gram-positive monoderms, with a single cell membrane and a thick peptidoglycan wall, and Gram-negative diderms with a thin peptidoglycan wall between two cell membranes[36], although many Bacteria have been shown to exhibit various atypical cell envelopes with a mixture of different characteristics[37]. It has often been suggested that having a double-membrane is a derived state[36], though more recent work has raised the possibility for a diderm ancestor[36,38]. The latter would be consistent with our phylogenetic analyses that resolve diderm Bacteria on either side of the possible roots. In agreement with this, we found many genes for lipopolysaccharide biosynthesis proteins, 24 of which had a PP>0.5 (8 with PP>0.9, see Supplementary Table 5), including the key proteins LpxC (K02535, PP=0.99/0.99/0.98) and KdsA (K01627, PP=0.92/0.9/0.78). In addition to lipopolysaccharides, two proteins used in transport across the outer membrane are recovered with PP>0.5, BamA (K07277, PP=0.65/0.61/0.29) and OmpH (K06142, PP=0.94/0.93/0.82). The inferred presence of these proteins in LBCA lends further support to the hypothesis that LBCA was a diderm, featuring an outer membrane with a full complement of lipopolysaccharides. We additionally recovered various genes encoding proteins for the construction of the cell wall and cell envelope, including 14 proteins predicted to be involved peptidoglycan biosynthesis had a PP>0.5 in roots 1 and 2 (Supplementary Tables 4 and 5).

Our analyses also suggest that LBCA possessed a fully functioning flagellum. More than half of the genes involved in flagellar construction are found with a PP>0.5 in all three roots, with 35/46 present in Roots 1 and 2 (see Supplementary Tables 4 and 5), including components of the C, Ms and P rings, the hook and hook-filament junction, as well as the Type III secretion system. We also recover three flagellar genes unique to diderm bacteria, flgH (PP=0.92/0.9/0.77), flgI (PP=1, 0.99, 0.99), which code for the L and P ring respectively, as we as flgA (PP=0.96/0.95/0.88). The corresponding proteins anchor flagella in diderm membranes, indicating that LBCA had a typical gram-negative flagellum and a double-membrane. In addition to a flagellum, we find 15 proteins for the construction of pili (PP>0.5), including 5 that seem implicated in the synthesis of a Type IV pilus. The key protein PliQ (K02666), which anchors the pilus in the outer membrane, is recovered with PP=0.86/0.82/0.62 across the different roots.

*Carbon metabolism, autotrophy and respiratory complexes*

The largest category of genes mapped to LBCA encode proteins involved in metabolism and transport of amino acids, coenzymes, nucleotides, inorganic ions and carbohydrates including pathways involved in central carbohydrate metabolism.

There are several carbon fixation pathways in extant autotrophic Bacteria including the Wood-Ljungdahl Pathway (WLP), the reverse TCA cycle, the Calvin cycle, and the 3-hydroxypropionate bicycle as well as distinct variants for the different pathways[39]. While the large subunit of the key enzyme of the Calvin cycle, i.e. Ribulose-bisphosphate carboxylase (RubisCO) and RubisCO-like proteins (RLP) are widespread in microbes and represent one of the most abundant protein families in the biosphere, we found only moderate support for the presence of a large subunit RubisCO-encoding gene in in LBCA (PP-range in the three roots: 0.24-0.59), in agreement with hypotheses in which the Calvin cycle and perhaps the carboxylation function of RubisCO/RLP evolved late[40].

In contrast, the reverse TCA cycle with the hallmark enzyme ATP citrate lyase, has been suggested as a possible ancient carbon fixation pathway[41–44]. While our analyses do not support the presence of ATP citrate lyase in LBCA, we do identify other enzymes of the TCA, including a citrate synthase as well as subunits of an Oxoacid:ferredoxin oxidoreductase, which may function in the TCA in both the oxidative and reductive direction (Extended Data Figure 7, Supplementary Table 5). For example, it has recently been shown that the citrate synthase, originally thought to operate only in the oxidative direction, can in fact catalyse the reverse reaction and allows to fix carbon in the facultatively chemolithoautotrophic thermophile Thermosulfidibacter takaii ABI70S6[44]. It has also been suggested that ATP citrate lyase may have emerged at a later stage from the domains of citrate synthase and succinyl-CoA synthase[44,45], which may further suggest that the TCA could operate in the reductive direction without ATP citrate lyase. Therefore, our analyses do not exclude the possibility that LBCA was capable of using the reverse TCA cycle to fix carbon.

Additionally, the WLP is generally thought to represent an ancient carbon fixation pathway on the basis of both biogeochemical and phylogenetic arguments[1,39,46,47] and previous phylogenetic work has suggested its presence in both the archaeal[1,47] and bacterial[47] common ancestors. While components of the methyl branch of the pathway were mapped to the root with PP support >0.95 for all roots, PPs for the subunits of the hallmark enzyme of the WLP, the Carbon monoxide dehydrogenase/acetyl-CoA synthase (CODH/ACS) were only moderate (i.e. PP >0.75) (Extended Data Figure 7, Supplementary Table 5). Considering the the methyl-branch of the WLP is also involved in alternative metabolisms including formate and folate transformations[48,49], it remains unclear whether the lack of strong support for a CODH/ACS in LBCA indicates that the WLP was absent in the bacterial ancestor or simply reflects the difficulty of mapping genes to the root with high statistical support. CODH/ACS subunits are not as widely distributed in extant Bacteria (Extended Data Figure 7, Supplementary Table 5) as in Archaea, so that their presence in LBCA would require extensive subsequent loss or HGT throughout the diversification of bacteria or suggests the later acquisition of this enzyme complex mediating the carbonyl-branch of the WLP.

On the other hand, our analyses provided strong support for the presence of Phosphate acetyltransferase and Acetate kinase, enzymes synthesizing acetate from acetyl-CoA (K13788, PP=0.86/0.9/0.74; K00925, PP=0.997/0.997/0.98)[50]. Furthermore, we find all six subunits of a $Na^+$-translocating ferredoxin:$NAD^+$ oxidoreductase (Rnf) complex, comprised of key genes *rnfA* (K03617, PP=0.99/0.99/0.95), *rnfB* (K03616, PP=0.84/0.95/0.70), *rnfC* (K03615, PP=0.77/0.89/0.59), *rnfD* (K03614, PP=0.89/0.94/0.78), *rnfE* (K03613, PP=0.89/0.96/0.77), and *rnfG* (K003612, PP=0.56/0.74/0.34) in LBCA. The Rnf complex is a membrane-bound respiratory enzyme that couples the oxidation of reduced ferredoxin ($Fd_{red}$) to the reduction of $NAD^+$ through a flavin-based electron transport chain, which is concomitantly coupled with the translocation of $Na^+$ ions, generating a transmembrane $Na^+$ motive force[51]. In the anaerobic acetogen, *Acetobacterium woodii*, this chemiosmotic gradient is used to drive ATP synthesis via a $Na^+$-dependent $F_1F_0$ ATP synthase [52]. Evidence has shown that the Rnf complex can function reversibly, where it catalyzes the reduction of oxidized ferredoxin with NADH using a chemiosmotic gradient ($H^+$/$Na^+$) generated via ATP hydrolysis [53,54]. Electron bifurcation through the redox coupling of NADH and ferredoxin allows for the production of high-energy intermediates from low-potential electron donors, which can be used to reduce $CO_2$ in the Wood Ljungdahl pathway (WLP)[55,56]. Taken together and in agreement with previous work suggesting the antiquity of the WLP[1,39,46,47], this leaves open the possibility for the presence of the WLP in LBCA and its capability of facultative acetogenic growth[50].

Our ancestral reconstructions indicated that LBCA encoded both membrane-bound F- and V-type ATP synthases. For the F-type ATP synthase, with recover all subunits PP>0.5 for roots 1 and 2, with the exception of subunit b. The alpha, beta, a and c subunits are recovered in all roots with PP >0.9. For the V-type ATP synthase, all subunits are found with PP >0.5 in all roots, except for subunits K and G/H (see Supplementary Table 5). Further, we identify the three subunits of a respiratory nitrate reductase (Nar)[57] with moderate PP values across the three root positions. NarH (K00371) has the highest support across the all three root positions (PP=0.84/0.81/0.6), with NarG (K00370) being recovered with moderate support in Roots 1 and 2 (PP=0.62/0.57/0.26) and NarI (K00374) only being recovered in Root 1 with moderate to low support (0.5/0.48/0.18). This suggests that LBCA might have had the ability for anaerobic respiration of nitrate.

Other components of the electron transport chain are patchily distributed across the different roots, with only a few subunits of each complex being found with PP >0.5 in all given roots. For instance, while difficult to annotate accurately in the absence of genome content and gene cluster information, we found subunits related to hydrogenases[58] (e.g. K18023, PP=0.96/0.97/0.89; K15830, PP=0.99/0.98/0.93), including a protein family comprising large subunits of [Ni-Fe]-hydrogenases (K00333, PP=0.80/0.92/0.58), in agreement with the hypothesis that hydrogen was a primordial electron donor[59,60]. However, we also find the subunits NuoG (K00336), CytB (K00412), and heme-copper type oxidase (CoxA) (K02274) in all three roots with PP >0.8 (Supplementary Table 5). Support for terminal oxidases are unexpected given that the atmosphere of the early Earth is predicted to be anoxic[61]. Interestingly, these genes were also recovered in a study that inferred the gene set present in the last universal common ancestor[62]. One possible explanation for these results is that aerobes are overrepresented among sequenced prokaryotic genomes; the wide distribution of these enzymes across the tips of the tree could then increase their probability to be mapped to the root in comparative analyses. Similarly, the relative patchy distribution of the key enzymes of the WLP across the tips of the bacterial tree could result in relatively low PPs (see above).

*Defense mechanisms, CRISPR*

Interestingly, we find several CRISPR-associated (Cas) proteins inferred to the root, suggesting the presence of a putative CRISPR-based prokaryotic immune system in LBCA. Highly-supported families (PP>0.95) belong to the Class 1 CRISPR-Cas systems, including the universal and essential Cas protein, Cas1 (K15342, PP=0.96/0.93/0.89), and three Cas proteins belonging to the Type III effector complex, Cas5 (K19139, PP=0.96/0.90/0.86), Cas7 (K19140, PP=0.96/0.90/0.85), and SS (K19138, PP=0.98/0.93/0.89). An additional nine Cas proteins belonging to Type I and III systems, were inferred to be present in LBCA with PP>0.80 (Supplementary Table 5). The presence of eight additional Cas proteins in the root was supported with PP>0.50. Here we find low support for Cas10 (K19076, PP=0.46/0.26/0), the signature cleavage protein of Type III systems. With the exception of Cas10, we recover all other essential and dispensable elements of a complete Type III system with weak to high support in our reconstructions suggesting the likely presence of this CRISPR system in LBCA.

The CRISPR-Cas proteins identified in this analysis were absent in the root of the CPR (Node 496) and rarely recovered across the taxa evaluated in this study (Supplementary Table 5, Figure 1), suggesting absence of CRISPR system components in members of the CPR clade. These findings are congruent with previous evidence showing that the CPR lack CRISPR-Cas systems possibly due to their host-associated or obligate-symbiont lifestyle[63]. Recent metagenomic analyses have uncovered two highly compact Class 2 CRISPR-Cas systems in uncultivated Bacteria, CRISPR-CasX and CRISPR-CasY, the latter of which was encoded in select CPR bacterial genomes[64]. We find no support/evidence for the key Cas proteins (CasX and CasY) of these novel CRISPR systems in our gene family reconstructions, suggesting a loss of the canonical CRISPR-Cas loci in this lineage and the later acquisitions of these novel systems in certain members of the CPR.

The majority of the CRISPR-Cas proteins recovered in our analysis belong to Class 1 systems, which exhibit greater architectural complexity and diversity in their effector modules compared to their Class 2 counterparts. For this reason, the Class 1 CRISPRs are rarely used for genome modification despite representing up to 90% of CRISPR-Cas systems[65]. It is postulated that this multiplex nature of Class 1 effector modules, specifically those of Type III systems, likely arose through a series of duplications and fusions of ancestral RNA recognition motifs (RRM)[66]. While the origin, organization, and composition of the CRISPR ancestor remains enigmatic, recent evidence has shown that a built-in signalling pathway in Type III systems comprised of nucleotide-binding (CRISPR-Associated Rossman Fold, CARF) and RNase (Higher-Eukaryote and Prokaryote Nucleotide-binding, HEPN) domains may be a key determinant between programmed cell death or induced dormancy and a targeted immune response[66–68].
